# Probing Individual Differences in the Topological Landscape of Naturalistic Brain Dynamics

**DOI:** 10.1101/2024.06.20.599966

**Authors:** Junxing Xian, Yini He, Yan Yan, Xiaohan Tian, Yingjie Peng, Jing Lou, Xiya Liu, Qi Wang, Tian Gao, Qi Wang, Yuqing Sun, Puze Li, Yue Wang, Shangzheng Huang, Kaixin Li, Ke Hu, Chaoyue Ding, Dazheng Li, Meng Wang, Bing Liu, Ang Li

## Abstract

Psychiatry seeks to unravel brain dysfunction and individual differences in real-world contexts. Naturalistic stimuli, like movie watching, are increasingly recognized for eliciting complex, context-dependent neural activity with high ecological validity. Yet, current methods often rely on standard paradigms that average data across time, limiting the full potential of such stimuli. Here, we present STIM, a Topological Data Analysis-based framework designed to dynamically track how individuals integrate complex contexts in real time. Applied to large-sample fMRI data from movie watching, STIM constructs a robust low-dimensional dynamical landscape that reflects group consensus while probing individual variations at both global (spanning narratives) and local (within specific narratives) levels. At the global level, individual differences emerge along a center-periphery gradient in the dynamical landscape, which significantly predicts fluid intelligence, underscoring the importance of neural adaptability and diversity. At finer scales, local geometric features correlate with context-specific psychological traits beyond cognition. STIM also captures developmental changes in the dynamical landscape and reveals abnormalities in conditions such as autism. These findings demonstrate that STIM leverages the rich information from movie stimuli and fMRI recordings as neural ‘probes’ to assess individual differences in cognition and mental health.

## Introduction

People live in highly complex social environments, where individual differences are largely reflected in how we perceive and respond to external stimuli. In extreme cases, such as mental illness, these processes can diverge significantly from those in healthy individuals, showing distinct cognitive, emotional, and attentional preferences.

Standard neuroscience experimental paradigms struggle to provide the rich stimuli characteristic of real-life conditions. Task paradigms, typically highly controlled, are designed to target specific brain circuits(*1*, *2*) related to particular cognitive or emotional functions. The variability in brain activation patterns between individuals can predict task performance and corresponding phenotypic traits(*3*). In contrast, the resting-state paradigm aims to reveal brain’s intrinsic functional organization, which has demonstrated stable individual specificity and can predict behavior and task-related activation across multiple domains(*4*). However, due to their highly tailored design, tasks often focus on specific functions and require repeated trials to accumulate brain activation patterns, potentially sacrificing generalizability for specificity. While the task-free resting-state paradigm offers broader applicability, it could reflect uncontrolled, internal mental states, which are difficult to measure and susceptible to interference. For instance, previous studies indicate that a significant proportion of individuals experience reduced alertness or even light sleep during resting-state scans(*5*, *6*).

The emerging naturalistic paradigm, using stimuli like movies, offers a more realistic and holistic approach to studying individual differences(*7*). This integrated paradigm combines the strengths of both task-based and resting-state paradigms, enhancing ecological validity and enabling high-throughput exploration of the brain’s complex functional systems(*8*).

However, current computational models for naturalistic stimuli lack the temporal resolution needed to capture how individual brains integrate context-dependent dynamic content(*9*).For instance, in a film scene with short, subtle negative cues, most healthy individuals may follow the main storyline without distraction, while individuals with depression might focus on the negative cues, leading to distinct subjective experiences. Notably, commonly used inter-subject correlation (ISC) and other similar approaches, measure the synchrony between different subjects over an entire duration or sliding time window(*10–15*), tend to blur transitions between brain states. State-based approaches such as the Hidden Markov Model (HMM)(*16*, *17*) and Co-activation Patterns (CAPs)(*18*) are constrained by the number of brain states they can account for (typically, n < 10), making them better suited for resting states and presenting challenges in accurately measuring the brain representations of rich stimuli.

Recent studies across species suggest that large-scale coordinated neural activity could map to continuous, latent space trajectories (or flows, manifolds) that supporting ongoing behavior and cognitive function at the system level(*19–22*). Inspired by these findings, we hypothesize that the fluctuating main storyline in a movie acts as an anchor or attractor basin guiding low-dimensional brain latent trajectories that may encode subjective experiences. These trajectories, likely stable at the group level, may serve as a neural signature of group consensus. Individual-specific experiences, on the other hand, would appear as divergent latent dynamics, shaped largely by individual characteristics; for instance, individuals with depressive tendencies may be more influenced by negative cues.

Here, we introduce STIM (Synchronized Topological Individual Mapper), a Topological Data Analysis (TDA)-based framework that aligns high-dimensional whole-brain activities across individuals during movie fMRI into a shared low-dimensional state space, while preserving flexible and unstructured individual brain dynamics. STIM generates a stable group-average dynamic landscape and quantifies individual-specific deviations from the group’s latent trajectories across two scales: a global, shape-like topology (spanning narratives) and a local, cluster-like geometry (within specific narratives). We expect STIM to ‘decode’ individual traits in a context-dependent manner, which can then be validated through self-reported cognitive and mental health phenotypes from questionnaires. Specifically, STIM uses the TDA-based Mapper approach to map high-dimensional datasets as a low-dimensional shape graph to visualize and analyze the topological and geometric information therein(*23*). Mapper has been shown to sensitively capture brain state transitions up to ∼4-9s faster than traditional methods, without requiring any annotation or assumptions about the analyzed data and can maximally retain the full spatiotemporal dynamic information at the individual level(*24*).

We applied the STIM framework to two large datasets: 170 subjects from the Human Connectome Project (HCP) and 970 subjects from the Healthy Brain Network (HBN), to examine how brain dynamics during movie watching relate to individual differences across multiple domains. We found that a stable group-level dynamical landscape characterizing individual differences could be constructed with as few as 20 participants. Both datasets indicated that global topology (spanning narratives) is significantly associated with cognitive ability, while local geometry (within specific narratives) explains specific dimensions of mental health. In most cases, individuals with higher cognitive abilities are more likely to follow the group’s latent dynamic patterns under more cognitively guided conditions. However, when negative cues within specific narratives are present, individuals with context-dependent mental health issues may deviate from group patterns. Based on the HBN dataset, we further explored the potential of the STIM framework in characterizing adolescent development and its applications to conditions such as autism.

## Results

### Mapper captures rapid and complex dynamics during movie-watching in the individual brain

In multi-task and resting conditions, Mapper is effective in constructing the individual dynamic landscape and capturing transitions between tasks as well as within spontaneous brain states(*25*). Extending previous studies, we expect that the TDA-based Mapper can effectively characterize the topology and dynamics of brain configurations related to more complex movie stimuli at the individual level, without necessitating any movie annotations.

Accordingly, we analysed high-quality 7T fMRI data during movie-viewing from the HCP (n=170, 4 runs for each participant), consisting of concatenated movies covering various themes such as science-fiction (sci-fi), drama, and nature scenes (details in Supplementary Tab. 1). Using the classic Mapper approach, we generated a topological landscape at single subject level(*26*) (Fig. 1a, See Methods). Classic Mapper aims to capture topological and geometric information, serving as a generalized notion of coordinatization for high-dimensional data (i.e., brain activity across fMRI scanning). The Mapper begins with a filter function *f* to generate a low-dimensional embedding, typically employing nonlinear dimensionality reduction techniques (such as UMAP(*27*), t-SNE(*28*)). This approach is crucial to preserve the ‘nearness’ of data, a desirable property in manifold learning that helps to mitigate the effects of the ‘curse of dimensionality’. The low-dimensional filtered data were then encapsulated into overlapping bins, generating a rough image of the data’s topology. Within each bin, partial clustering was applied to refine the scale, with each cluster forming a node and the overlaps between nodes forming edges. This process produces a multiresolution image of the data’s topology and is less sensitive to the choice of metric. Through this processing, the high-dimensional brain activity data are condensed into a shape graph(*23*).

**Fig 1.**
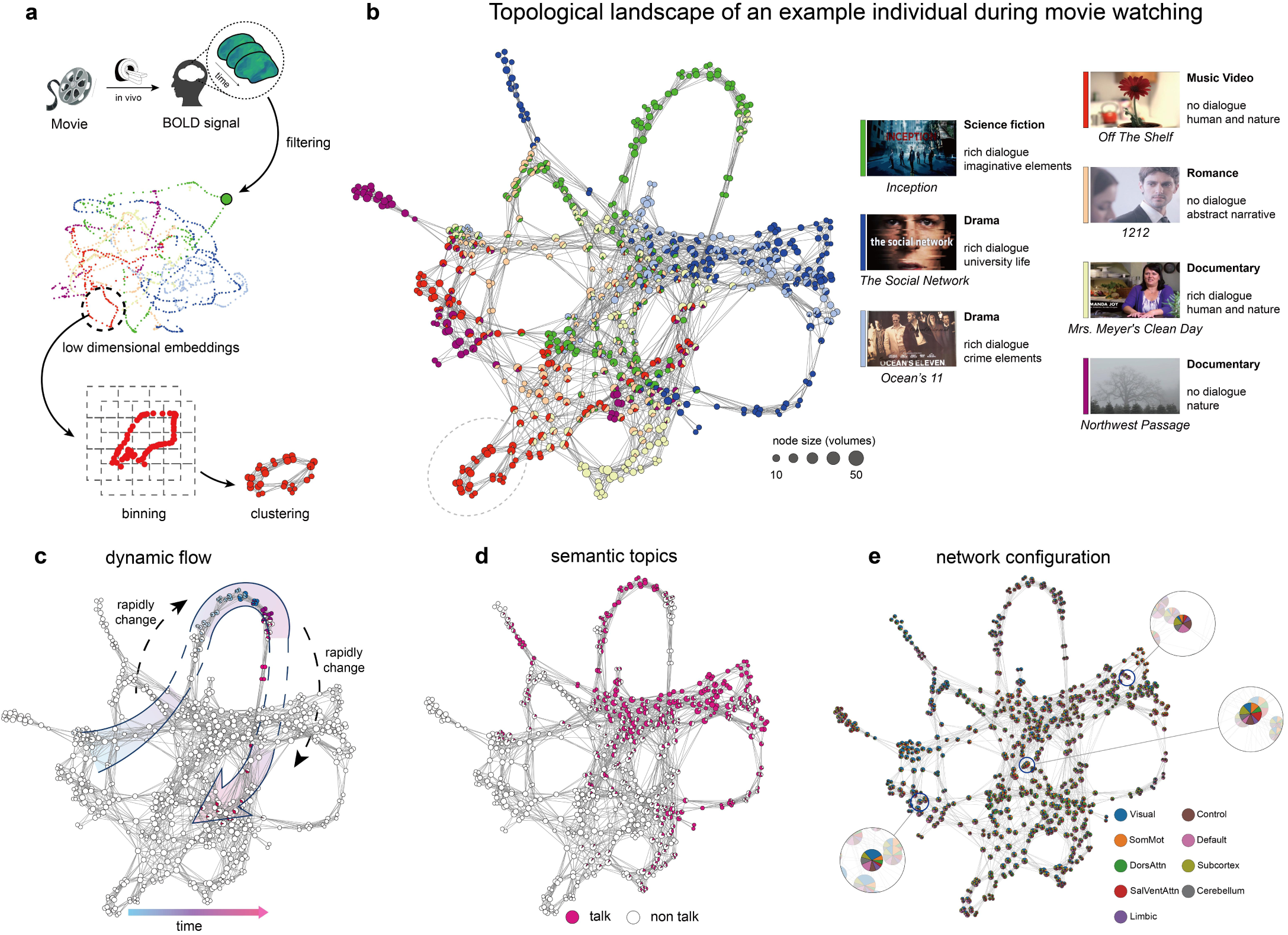
Mapper approach captures the individual dynamic topological landscape. **(a)** A conventional Mapper pipeline for movie-viewing fMRI analysis is outlined as follows: First, the high-dimensional neuroimaging data are embedded into a low-dimensional space using a nonlinear filter function f. Here, we chose Uniform Manifold Approximation and Projection (UMAP). Second, the low-dimensional embedding is mapped (via binning and clustering) into an overlapping grid, generating a shape graph, where the nodes of the shape graph are determined by bins, and the edges are determined by the overlap between bins. Within the shape graph, closely connected nodes correspond to highly similar whole-brain time frames. Lastly, annotations of interest are added to the shape graph at the volume scale (for example, time flow) or the node scale (for example, node centrality) to further research the topological relationships among different features and to facilitate their visualization. **(b)** The HCP dataset includes various categories of movie stimuli, and 7 of 18 representative clips were used as examples, including sci-fi (*Inception*), drama (*Ocean’s Eleven, The Social Network*), music video with nature scenes (*Off the Shelf*), romance (1212), documentary (*Mrs. Meyer’s Clean Day*), and nature scenes (*Northwest Passage*) (from left to right). **(c, d, e)** Three kinds of Mapper independent information were annotated for illustration, including time order (70 volumes), ‘talk’ semantic label, activation of nine pre-parcellated brain networks.

Fig.1b displays the topological landscape of global brain dynamics for a representative individual (see more people in Supplementary Fig.1), that is, a Mapper-generated shape graph. This graph intuitively illustrates nodes as clusters of whole-brain volumes with high similarity (with size indicating the number of included time volumes, from 4 to 48 seconds for each node), and nodes sharing overlapping volumes are connected. A distinctive feature of the Mapper method is its embedding of temporal brain activations into the topological structure of the graph, enabling a dynamic, global perspective analysis and visualization of high-dimensional fMRI data. A video version of Fig.1b, available in Supplementary Mov.1, concurrently showcases the dynamic trajectory of the shape graph and the corresponding movie content being watched. The color annotations in Fig.1b represent different movie sources. Interestingly, movies viewed in different runs can form a cohesive topological landscape, and movies with similar themes tend to exhibit stronger connections, such as dramas (*The Social Network* and *Ocean’s 11*) and nature scenes (*Off the Shelf* and *Northwest Passage*), while a theme-rich movie (*Mrs. Meyer’s Clean Day*) is dispersed throughout the graph. This suggests that merging multiple movies can establish a global, rich dynamic landscape.

Moreover, Mapper can provide an interactive, flexible visualization to interpret how the brain traverses its dynamical landscape during movie-watching, by annotating the shape graph with different types of information: i) When annotated by time order, it can display the flow and rapidly changing dynamics related to the viewing context (Fig.1c and Supplementary Mov.1); ii) When annotated by focused semantic topics, such as volumes with ‘talk’ or without, emerging clustering in the individual shape graph becomes visible (Fig.1d); iii) When annotated by network activation, the pie-chart notation of nodes represents the weight of functional networks, revealing interested brain configuration patterns(*25*) (Fig.1e).

### STIM framework to capture individual divergence in global topology and local geometry

While the Mapper approach effectively delineates individual dynamical landscapes, significant challenges persisted in characterizing the neural signature of consensus at the group level and the divergence specific to individuals. Also, previous studies focused on non-event-related topological descriptors cannot adequately accommodate analysis of specific movie content(*16*, *29*, *30*). Therefore, we introduce the STIM (Synchronized Topological Individual Mapper) framework, aiming to construct a group-level dynamical map of consensus as the reference and quantify individual-specific divergence both in the global topology, corresponding to the overall viewing experience, and in local geometry, corresponding to the experience of specific plot elements. STIM presupposes that individual-specific low-dimensional, latent dynamics, encoded by the Mapper shape graph, reflect subjective experiences during movie viewing, wherein the same stimuli evoke consensus or divergence across viewers. Thus, STIM was developed to align individuals while preserving and quantifying individual-specific topological features.

For a more intuitive demonstration, we chose the segment from the movie *Inception* (Fig. 2a, duration: 3 minutes and 46 seconds) as an example, and manually annotated two scene transitions, both of which feature a significant change from face close-up/dialogue to sci-fi/space folding content (the two sci-fi/space folding scenes are named ‘city-folding’ and ‘mirror corridor’). Fig. 2b shows shape graphs of three participants watching *Inception*, with annotations of movie plots (light green and light purple represent the first close-up/dialogue and ‘city-folding’, connected by transition-A; deep green and deep purple represent the second close-up/dialogue and ‘mirror corridor’, connected by transition-B). In the low-dimensional topological space, two typical individuals (subject-1 and subject-2 in Fig. 2b) were observed to exhibit dynamical consensus patterns: integrating into two different clusters associated with face close-up/dialogue and sci-fi/space folding, respectively, and exhibited an ‘expected’ dynamic transition from the face close-up/dialogue topological region to the sci-fi/space folding topological region. Interestingly, some individuals exhibited ‘unexpected’ transitions towards two distinct topological regions, potentially indicative of their divergence in movie-viewing experiences (such as subject-3 in Fig.2b). The shared and specific features were plausibly observed in individual shape graphs (Fig.2b). However, the shape graphs were independent and thus incomparable. To construct a shared topological space, we concatenated two individuals’ data to provide a higher-dimensional representation space and built a combinatorial shape graph of subject-1 and subject-3 (Fig. 2c left), which can represent the synchrony and idiosynchrony: their dynamics cluster at the two face close-up/dialogue and the ‘mirror corridor’; however, subject-3 deviated to another topological region while watching the ‘city-folding’ after transition-A (A full video version is available in Supplementary Mov.2). Therefore, an important procedure in STIM is to concatenate the high-dimensional brain activities across subjects to align brain dynamics and generate a shared low-dimensional topological space using Mapper (see Fig. 2c right). Furthermore, the shared shape graph can be converted into an adjacency matrix (i.e., temporal connectivity matrix, TCM)(*24*), in which elements represent the connectivity strength between the two volumes by shared nodes. By calculating the TCM of the combinatorial shape graph, we can quantify the topological similarity between any volumes from different individuals (see Methods for details).

**Fig 2.**
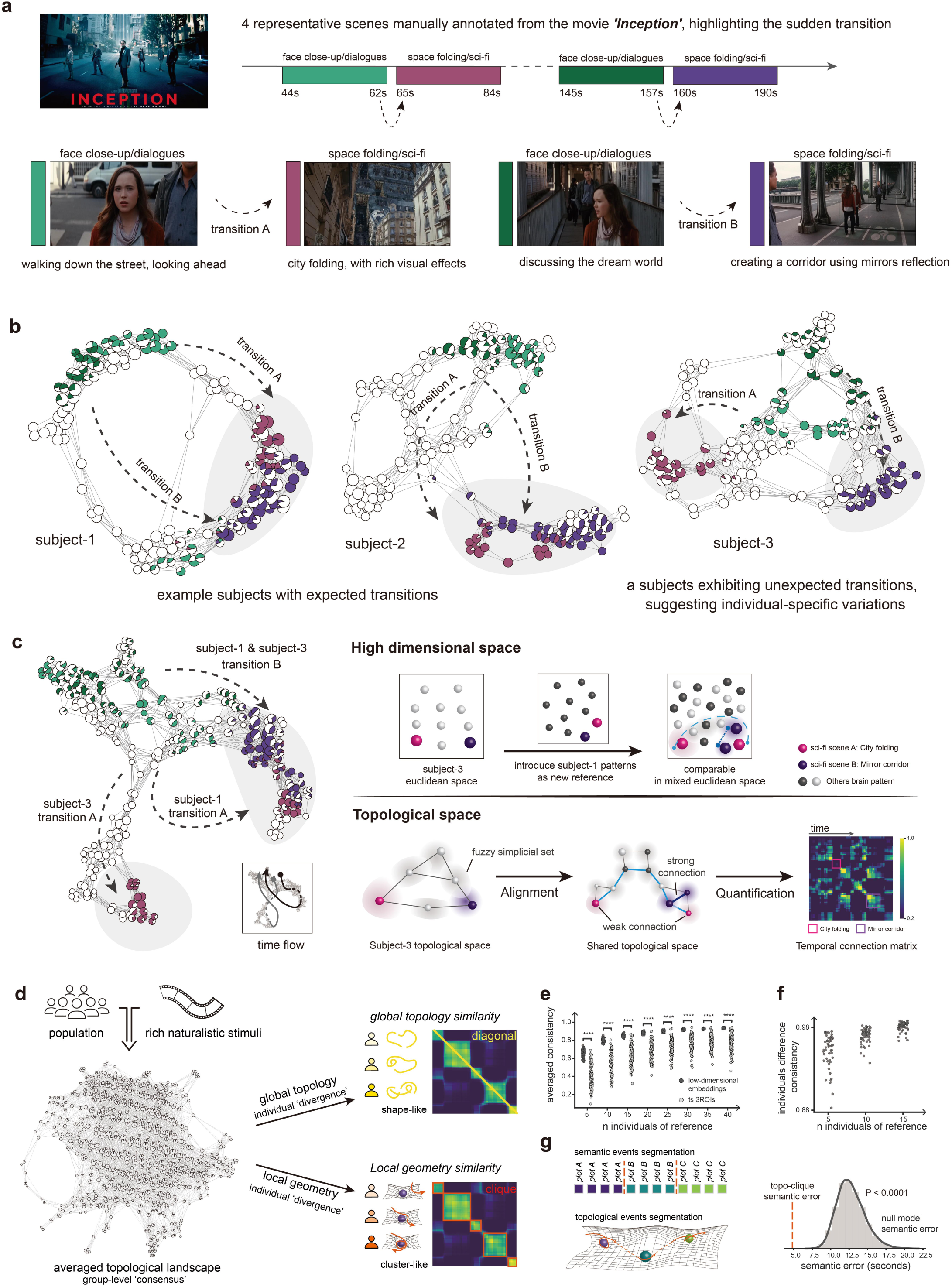
The STIM (Synchronized Topological Individualized Mapper) framework. **(a)** We manually labeled two representative segments from Inception, each with a continuous sequence, featuring face close-up/dialogue in the first half and sci-fi/space folding scenes in the second half. **(b)** We annotated four scenes in the individual shape graphs of the three subjects chosen as examples. According to our assumption, brain dynamics are clustered during face close-up/dialogue scenes and sci-fi/space folding scenes at different times. Subject-1 and Subject-2, conforming to the assumption, respectively formed different clusters, while Subject-3 exhibited an “unexpected” transition. **(c)** STIM framework. The left panel shows the shape graph of alignment for Subject-1 and Subject-3, with annotations of the four scenes. The right panel illustrates the motivation for alignment: introducing a new subject in high-dimensional Euclidean space to provide a comparable combination of TRs. In the topological space, the distances between volumes from different individuals are rescaled through a fuzzy simplicial set (in the context of UMAP). Subsequently, through the filtering and binning steps, a combinatorial shape graph (aligned shape graph) is generated. Finally, the strength between volumes is quantified by TCM. A time-flow arrow illustrates the temporal dynamics of the two subjects. **(d)** Two scales of topological similarity. Global topological similarity is defined as diagonal of subject-group interact TCM (yellow line), represents the overall shape similarity. Local topological similarity is defined as weighted sum of elements in a cluster (orange box), represents deviation in specific movie content. **(e)** Group consistency across different sample sizes between STIM and time series: the scatter plot displays the Pearson correlation between two family unrelated groups with increasing sample sizes, ranging from 5 to 40 subjects, randomly selected. For a fair comparison, each trial randomly selects time series from 3 ROIs. ****: P<0.0001 **(f)** Stability test of global topology similarity. We changed the group reference by randomly selecting different small group of individuals. We examined the stability of global topology similarity across different group sizes. Scatter plots show the Pearson correlation of global topology similarity between small group references and the whole group reference. **(g)** An illustration of the ecological validity of topologically self-organized cliques: The dashed line indicates the error between the topological events segmentation and the semantic events segmentation, accompanied by a gray null distribution, repeated 10,000 times with 161 events segmentations randomly divided in each iteration.

The STIM framework is designed to effectively analyze large-sample movie-viewing data. To characterize the canonical dynamics as group-level consensus, STIM concatenated the volumes of all movies and all individuals (for HCP: 2804 volumes×170 subjects, 476,680 volumes) to create a panoramic view of movie-viewing dynamics. The ‘shared filtering’ data were subsequently binned and clustered to derive each subject’s TCM relative to the group reference, which was generated by averaging embeddings across individuals in a low-dimensional space (Fig. 2d). STIM then quantified individual divergence by decomposing the individual-group TCM into two distinct scales (Fig. 2d, see Methods):

i. *Global topology similarity*: This metric quantifies the overall shape similarity of topological landscapes cut across movie contents. As an example, Subject-3 might not only deviate in the ‘city-folding’ scene of *Inception* but also exhibit lower similarity with the group reference across various movie scenes, indicating a lower global topology similarity, reflecting broader deviation in brain dynamics under different cinematic contexts. Global topology similarity is defined as the diagonal value of the TCM (n volumes = 2804 for all HCP contents). STIM filtered low-dimensional embeddings (or trajectories) showed a more consistent population-shared shape of global topology than the original time series (Fig. 2e). Furthermore, trajectories of latent dynamics in the STIM pipeline showed significantly higher consistency than top region of interests (ROIs) within the visual network (Supplementary Fig. 2), supporting that the latent dynamics of consensus reflects a global integration of the brain during movie-watching beyond merely sensory input. This global integration may be linked to subjective experience. For example, in *Off the Shelf*, the first half presents unguided, non-narrative natural scenes, followed by a clearly structured story segment. Group clustering associated with the narrative appeared in low-dimensional space, rather than in visual regions, potentially reflecting shared subjective experiences (Supplementary Mov.3 and Supplementary Fig.3). Notably, the individual differences in global topology similarity remained stable even with a small reference sample. For example, when randomly selecting 15 subjects as the group reference, the average Pearson correlation *r* across iterations was 0.98 (Fig. 2f). We also constructed a shared topological dynamic space for *Inception*, where different groups (with *n* = 15 per group) exhibited very highly similar transitions and positions over time within the topological space (Supplementary Mov.4).
ii. *Local geometry similarity*: This metric quantifies the local similarity elicited by specific movie contents. For example, Subject-1 and Subject-3 showed lower similarity in ‘city folding’ content, while forming a cluster-like structure in the second sci-fi scene (Fig. 2c, ‘mirror corridor’ in Fig. 2a). Local geometry similarity is defined as the weighted score of cliques detected by a change point detection algorithm(*31*) (n clusters = 161 across all HCP movie contents, see Methods). To ensure that the algorithm can capture meaningful movie transitions, we compared these topological cliques to the manual segmentations based on semantic labels manually annotated by the HCP dataset (see Methods). We found that the discrepancy between topological and semantic cliques was significantly smaller compared to a null model (*P* < 0.0001, Fig. 2g), indicating that local topological similarity can capture fine-grained, unstructured transitions in movie plots. The deviation between manually labeled transitions (A and B in *Inception*) and the detected change points is shown in Supplementary Fig. 4.

### The principal gradient of individual differences in the topological landscape explains cognition from a dual perspective

Global topology similarity can capture the individual divergence in the shape of the dynamical landscape. We next explore how this divergence in global topology differs between individuals and over time by integrating multi-level information, including topological attributes within the landscape, synchronous modes within the population, brain network configuration, movie contents, and behavioral domains. We performed a principal component analysis (PCA) on the global topological similarity matrix (170 subjects×2804 volumes) across time.

Fig.3a highlights the top 5% positive and negative volumes of the weights in the first principal component (PC1) of the group shape graph, denoted as +VOLs and -VOLs (140 volumes), respectively. At a global level, -VOLs and +VOLs exhibit a trend from the graph’s center to its periphery. Further, employing betweenness centrality to measure topological attributes(*32*), we discovered that + VOLs exhibited significantly lower betweenness centrality than - VOLs (t=-13.9, *P* < 0.0001, Fig.3a).

Besides topological attributes, we examined how the + VOLs and - VOLs systematically differed in group synchronous modes. We observed a significant divergence in the dynamics of synchrony between +VOLs and -VOLs (t=18.1, P<0.0001, Fig.3b). We presented the low-dimensional embeddings of all subjects’ volumes at two representative volumes for +VOLs and -VOLs (Fig. 3b). +VOLs exhibited an ‘attractor-like’ mode, where subjects tend to converge (stronger consensus); in contrast, -VOLs displayed an ‘unconstrained’ mode, where subjects are more dispersed (weaker consensus). These two modes also exhibited distinct preferences for movie content. We showcased the top 10 semantic labels where +VOLs and -VOLs differ most significantly in their relative proportions (Fig. 3c). We found that +VOLs were more associated with the human-related scenes, predominantly featuring the top entries such as ‘man’, ‘talk’, and ‘sit’. In contrast, - VOLs were more aligned with nature scenes compared to +VOLs, with the top labels being ‘forest’, ‘fog’, and ‘flower’.

**Fig 3.**
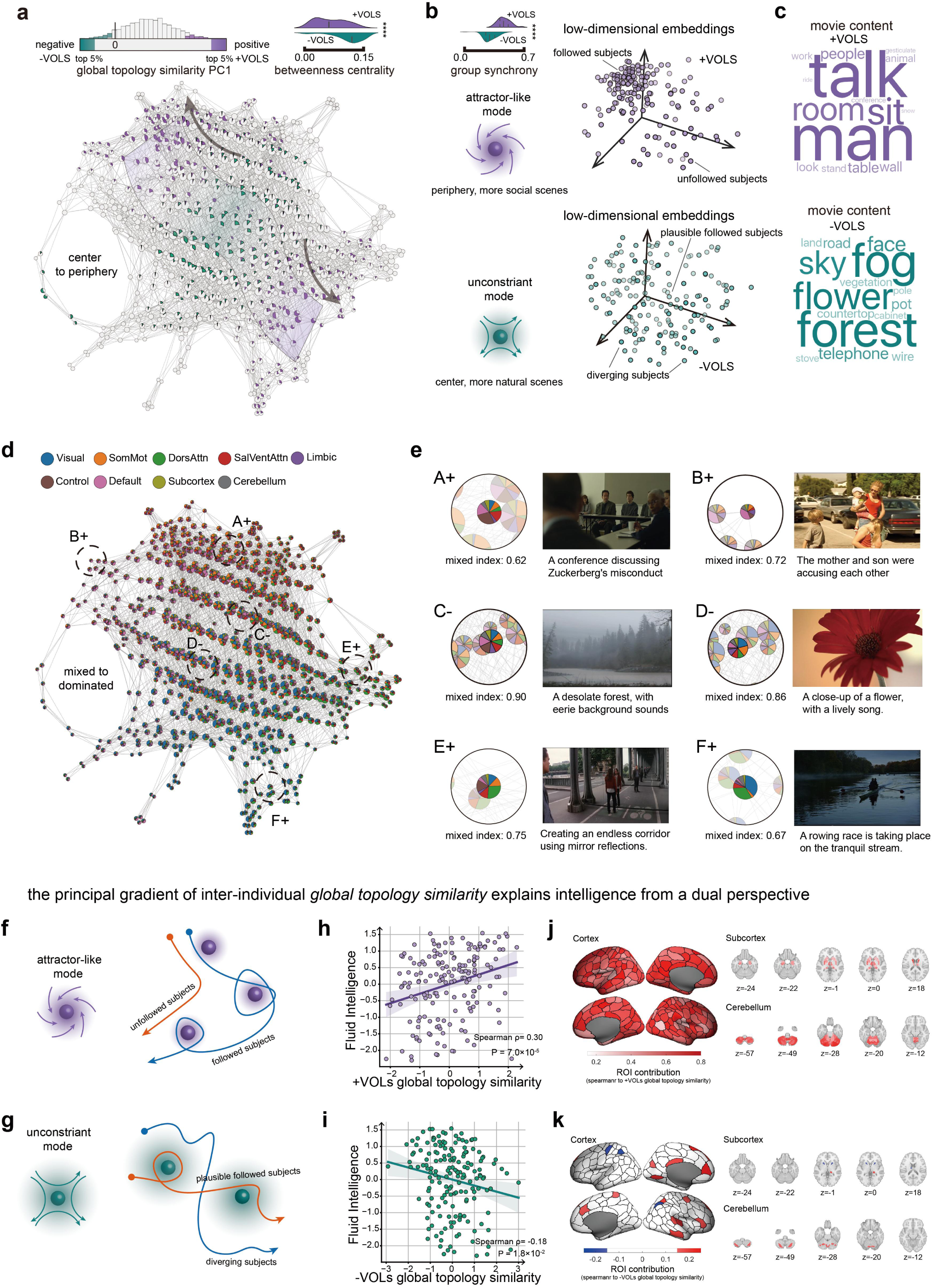
The principal gradient of individual differences in the global topology of the dynamical landscape. **(a)** We displayed a group shape graph from averaged embeddings and annotated time points at the extremes of PC1 (+VOLs and -VOLs). The distribution plot shows the betweenness centrality for +VOLs and -VOLs. **(b)** Population dynamics of +VOLs and -VOLs. The distribution plot illustrates the group synchrony for +VOLs and -VOLs. Group synchrony is defined as the average global topology similarity among individuals, corresponding to the illustration of attractor-like and unconstrained modes. UMAP embeddings for all individuals at the most extreme volumes are displayed. **(c)** Word clouds depict the 15 terms with the largest semantic frequency differences between +VOLs and - VOLs. **(d)** Brain network configurations are annotated on the group shape graph, with proportions representing network activation levels. **(e)** Across the most extreme time points in +VOLs and -VOLs, we displayed compressed information of six representative nodes, including network configurations, the mixed index, and corresponding movie content. **(f, g)** We illustrated different individual dynamic patterns in +VOLs and -VOLs, where orange lines represent individuals not following the group, and blue lines represent those following the group. **(h, i)** Scatter plots show the correlation between averaged global topology similarity and fluid intelligence for +VOLs and -VOLs. **(j, k)** The values demonstrate the contribution of ROIs, with the sign indicating the direction of correlation between ROI and PC1 of global topology. DorsAttn, dorsal attention network; SomMot, somatomotor network; SalVentAttn, salience/ventral attention network.

How is the principal gradient interpreted from the perspective of brain network configuration? Previous work showed a dynamical topographic gradient of resting-state networks(*25*). Similarly, we annotated the activation of 9 networks on the group shape graph and qualitatively depicted the network’s domination using pie-chart proportions (Fig. 3d). We calculated a mixed index of nodes to describe the overall activation variation across the networks (see Methods). We found that the mixed index for +VOLs was significantly lower than that for -VOLs (group-level across time: *t*=-11.35, *P* < 0.0001; subject-level across individuals: *t*=-5.27, *P* < 0.0001), suggesting that the network configuration along the principal gradient shifts from uniform activation to the dominance of one or more networks. Integrating movie content into our analysis, we highlighted six movie segments aligned with the principal gradient, highlighted by magnified nodes. Four of these segments belonged to +VOLs (A+, B+, E+, F+ in Fig. 3e) and were associated with social interactions or human behaviors, engaging networks such as the control network, default mode network, dorsal attention network, and visual network. The remaining two segments belonged to -VOLs (C-, D- in Fig. 3e), which were linked to non-human scenes.

Therefore, individual divergence in group consensus can be interpreted as follows: At the +VOLs, subjects engage in ‘attractor-like’ modes, where higher scores indicate greater flexibility in adapting brain configurations to the diverse cognitive contents (Fig. 3f). At the -VOLs, subjects exhibit ‘unconstrained’ modes linked to nature scenes and mixed network activations, with higher scores suggesting less mind wandering and greater conformity with the group (Fig. 3g). Interestingly, within +VOLs, individuals who align more closely with the group consensus (demonstrating higher global topology similarity) have greater fluid intelligence (Spearman r = 0.30, *P* < 0.0001, Fig.3h); in contrast, within -VOLs, individuals who diverge from the group consensus (exhibiting lower global topology similarity) display increased fluid intelligence (Spearman r = -0.18, *P* = 0.017, Fig.3i). The results suggest that the principal gradient of individual differences in global topology explains cognition from a dual perspective: in ‘attractor-like’ modes, stronger dynamic ‘following’ to group latent states correlates with better cognition; in ‘unconstrained’ modes, greater divergence correlates with better cognition.

Next, we calculated a global principal gradient across all volumes (including +VOLs and -VOLs) for each participant. This gradient is significantly associated with cognitive domains but not with emotional or personality domains (Supplementary Tab. 3). Fluid intelligence showed the strongest correlation (Spearman r = 0.32, P < 0.0001). This effect is comparable to the predictive capacity of static functional connectivity (sFC, Supplementary Fig.5). To ensure the gradient score as a trait-like marker, we computed principal gradients across two independent days, resulting in a relatively high correlation (Spearman r = 0.54, *P* < 0.0001). Lastly, we examined the contributions of the principal gradient to fluid intelligence at the brain region level in the population weights. In brief, we calculated the similarity between each ROI and the global topology across individuals (see Methods for more details). We found that +VOLs were associated with most brain regions, spanning the major cortical areas and cerebellum (Fig.3j), while significantly fewer regions correlated with - VOLs, involving the default mode network and the control network (Fig.3k).

### Local geometry similarity of individual differences in topological landscape explains mental health

We hypothesize that the local geometry similarity corresponding to topological cliques can explain multi-dimensional human behaviors in a manner related to the specific movie content. To explore the associations, we included 59 behavioral measurements, including HCP-defined domains in *Cognition*, *Emotion*, *Psychiatric and Life Function* (Fig. 4a, Supplementary Tab.2). Based on the STIM framework, we derived 161 modular topological cliques (average duration = 17.4 seconds) as a range of non-overlapped ‘attractors’ that organized by brain dynamics and movie contents (Fig. 4b). We anticipated that some of the cliques, particularly those rich in cognitive, emotional, and well-being content, could act as neural ‘probes’ reflecting corresponding domains of human behavior (Fig. 4c).

**Fig 4.**
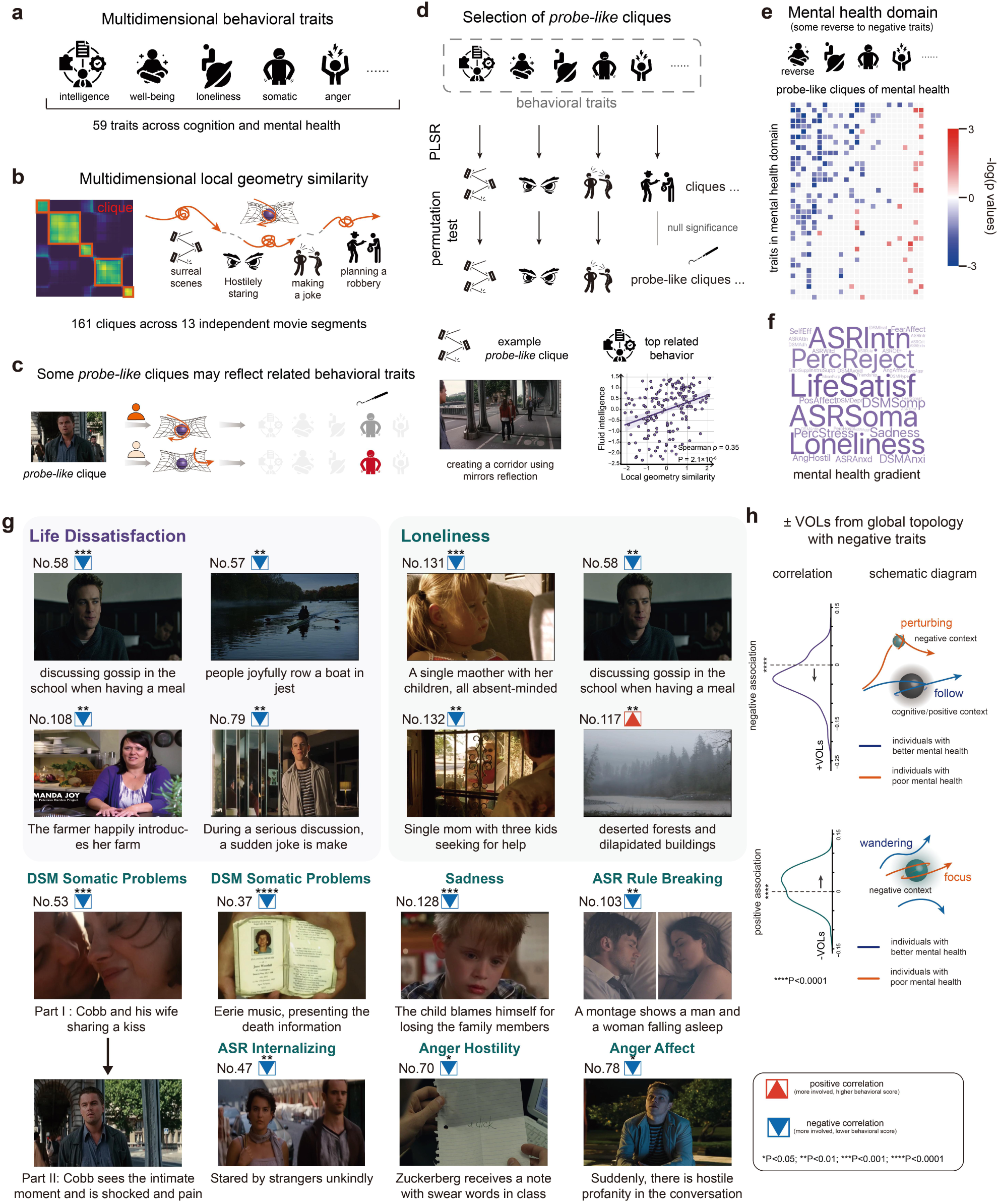
The individual differences in the local geometry similarity of the dynamical landscape. **(a)** Multi-dimensional behavior traits used for analysis. A total of 59 behavior traits were included. **(b)** Multi-dimensional local geometry similarity for 161 cliques, including various movie content. **(c)** Hypothesis of probe-like cliques: the individual local geometry similarity of specific clique may relate to corresponding behavior traits. **(d)** Methodological illustration for identifying probe-like cliques: PLSR with permutation test was conducted to identify probe-like cliques. An example of a probe-like clique related to fluid intelligence: characters create a surreal corridor using mirror reflection. **(e)** Subsequent analysis focused on negative mental health (some behavior traits were reversed). A heatmap displays significant correlation levels between probe cliques and negative mental health dimensions, arranged by mean significance. Non-significant points (P > 0.1) are shown in grey. **(f)** A weighted word cloud depicting the significance level for various negative mental health. **(g)** Movie contents corresponding to representative cliques significantly associated with mental health dimensions are presented. The selection includes the four most significantly correlated cliques with *loneliness* and *life dissatisfaction*, along with other representative cliques across all probe-like cliques and mental health dimensions (excluding *loneliness* and *life dissatisfaction*). Cliques No.108, No.57, No.128, and No.78 did not pass the permutation test; nevertheless, we included these pairs because of their interpretive value. P-values denote the raw Spearman correlations between clique-behavior pairs. The brain configurations of the displayed cliques are shown in Supplementary Fig.6. **(h)** Dynamic modes of group dynamics in negative emotions. The distribution diagram illustrates the direction of Spearman correlation between the topological similarity in +VOLs and -VOLs (i.e., ‘attractor-like’ and ‘unconstrained’ mode in Fig. 3) and individual top 10 negative emotions score. The schematic displays the population dynamics of the +VOLs and -VOLs with negative emotions.

We conducted a partial least squares regression (PLSR) to identify such ‘probe-like’ cliques which are behavioral relevant, resulting in significant 25 cliques (*P* < 0.05, after permutation). We next used pairwise correlation analysis to link specific behavioral domains to these ‘probe-like’ cliques (Fig. 4d, see Method). Using fluid intelligence as an example, a few topological cliques show significant correlations, especially ‘mirror corridor’, which had a comparable effect size to the global principal gradient (Spearman *r*=0.35, *P*<0.0001, Fig.4d). The *Cognition* domain was discussed in the previous section. Here, we focus on *Emotion* and *Psychiatric and Life Function*, combined as the mental health domain. Some measurements were reversed to align as the uniform negative traits for mental health, such as transforming *life satisfaction* to *life dissatisfaction* (see Methods). The local geometry similarity of most cliques negatively correlated with negative traits, while in a few cliques was positively correlated with the negative traits (Fig. 4e, significance denoted in the legend, only displaying pairs that are at least marginally significant: *P* < 0.1). The contribution of different mental health dimensions was visualized using a word cloud, where font size linearly represents the mean significance rank (Fig.4f).

Among these, the most relevant behavior is *life dissatisfaction*, for which we highlighted the top four most correlated cliques (Fig.4g).: joyfully discussing gossip while dining with friends (No.58, *ρ* =-0.29, *P*=0.00012); leisurely rowing a boat (No.57, *ρ*= -0.24, *P* = 0.0013); a lady happily introducing her farm accompanied by light music (No.108, *ρ*= -0.24, *P* = 0.0016); making a joke with a smile during a serious discussion (No.79, *ρ*=-0.23, *P*=0.0028). These scenes, accompanied by joyful facial expressions or relaxed conversational atmospheres, are positive aspects of life and social interactions. The second correlated dimension is *loneliness*, with the top four related cliques being: a single mother with her children remains silent, their expressions sorrowful, accompanied by sad music (No.131, *ρ*=-0.25, *P*=0.00099); the next clique, this single mother with three children seeks help (No.132, *ρ*=-0.22, *P*=0.0043); in a somber musical setting, deserted forests and dilapidated buildings are displayed (No.117, *ρ*=+0.21, *P*=0.0063) as well as No.58 (*ρ*=-0.21, *P*=0.0063). The scenes associated with loneliness almost invariably involved somber atmospheres and unhappy expressions.

The findings on *life dissatisfaction* and *loneliness* indicate that local brain geometry accounts for individual differences in areas related to specific, explainable movie content. We next identified cliques most closely associated with negative traits of mental health, with the representative examples displayed below (Fig.4g; see Methods; additional details in Supplementary Tab. 4): For *DSM (Diagnostic and Statistical Manual of Mental Disorders) Somatic Complaints*, the most related scenes: In the process of Cobb seeing himself being intimate with his wife, his facial expressions vividly convey the astonishment and pain of losing her (No.53, *ρ*=-0.25, *P*=0.00092); accompanied by somber music, an interviewee solemnly shows a photo of his deceased mother from his pocket (No.37, *ρ*= -0.32, *P*<0.0001). Both scenes involve intense emotional trauma related to death, which can elevate psychological distress to physical discomfort(*33*). Additional cliques-mental health pairs include: being continuously stared at unkindly by strangers, explaining interindividual *ASR (Adult Self Report) Internalizing Score* (No.47, *ρ*=-0.24, *P*=0.0021); viewing a child blames himself with a negative face, relating to *Sadness* (No.128, *ρ*=-0.25, *P*=0.00088); a montage sequence shows a man and a woman each lying on their sides in bed, corresponding to the *ASR Rule Breaking Behavior Score* (No.103, *ρ*= -0.22, *P*=0.0040). Exposure to hostile expression tends to relate anger dimension: the main character receives a note with ‘udick’ (*Anger Hostility*, No.70, *ρ*= -0.18, *P*=0.021), and a ‘damn’ word is suddenly used in a dialogue, provoking an angry response from another character (*Anger Affect*, No.78, *ρ*= -0.19, *P*=0.013).

Interestingly, the number of cliques negatively correlated with negative mental health significantly exceeds those with a positive correlation (Fig. 4e), suggesting that negative traits primarily disrupt individuals’ latent trajectories toward group consensus during specific movie scenes. We hypothesize that a dual perspective may also exist in the mental health domain: in a few ‘unconstrained’ modes cliques (such as No.117, the deserted forests), negative traits may prevent individual divergence, leading to immersion in a negative emotion. To verify this, we utilized +VOLs and -VOLs from the previous section, analyzing the directional correlation between individual differences and negative emotions (Fig. 4h). We found that individual differences in +VOLs are generally negatively correlated with negative traits (*t*=-8.8, *P* < 0.0001), whereas -VOLs are positively correlated with negative traits (*t*=4.7, *P* < 0.0001). This suggests that distinct individual dynamic patterns are linked to negative mental health in ‘attractor-like’ or ‘unconstrained’ modes (Fig. 4h).

### Using the STIM framework to characterize developmental progress and conditions

We further investigated whether the STIM framework could offer new insights into the developmental progress and conditions of brain dynamics. To this end, we utilized the Healthy Brain Network (HBN) dataset, which includes fMRI data from 970 participants aged 5-21, who watched a 10-minute clip of the animated film *Despicable Me*. This comedic segment contains numerous instances of social processing. The participants, presenting a wide array of transdiagnostic conditions (e.g., autism spectrum disorder), provide a comprehensive context for investigating developmental trajectories and the spectra of mental disorders (Fig. 5a).

**Fig. 5.**
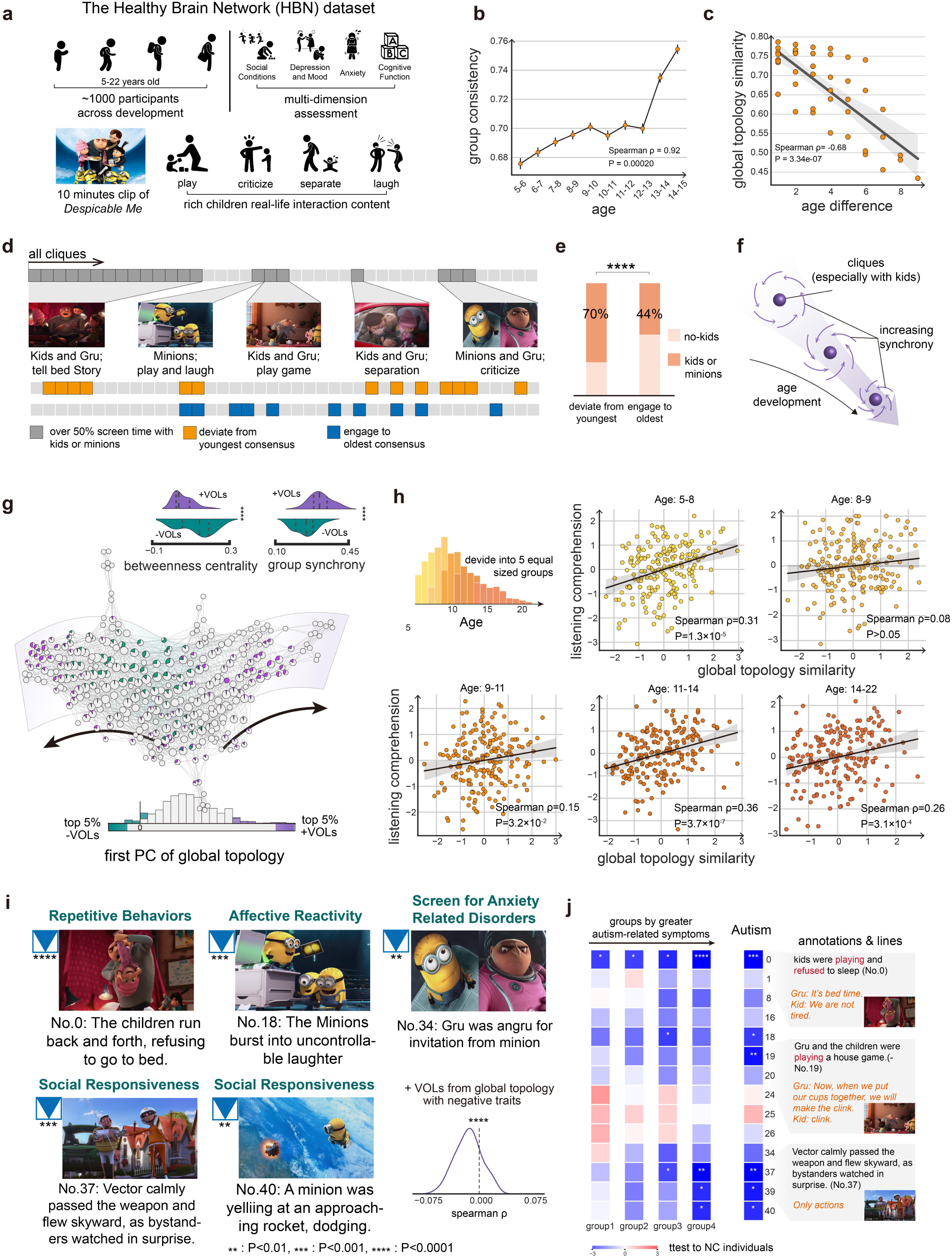
Application in developmental progress and conditions. **(a)** Overview of HBN dataset. **(b)** Point plot showing group consistency across different ages, indicated by Spearman’s correlation coefficient, with 95% confidence intervals shown. **(c)** Scatter plot of age intervals and group global topology similarity. **(d)** Gray barcode plots display cliques where ‘kids’ or ‘minions’ appear for over 50% of the screen time, with key characters and times described through snapshots and text. Orange barcode plots shows cliques that significantly deviate from the youngest consensus with increasing age. Blue barcode plots depict cliques significantly engaging into the oldest consensus as age increases. **(e)** Ratio of child appearances in cliques shows significant differences in younger individuals and older individuals. **(f)** An illustration of developmental characteristics of topological properties. **(g)** Shape graph of averaging individuals, with annotations of top 5% and bottom 5% time points of principal gradient. Violin plots show differences of two parts of time points in betweenness centrality and group synchrony (mean global topology similarity across individuals). **(h)** The study population is divided into five age groups, with scatter plots showing the relationship between age and listening comprehension and topological similarity. **(i)** Five representative cliques with description and corresponding mental health. Distribution plot shows the direction of top 5% timepoints in negative mental health. **(j)** Barcode plots show relationship between all probe-cliques and ASD (symptoms gradient and diagnostic assessment), with description of top three probe-like cliques.

We first focused on the application of STIM in developmental progress. To mitigate the impact of the age distribution of HBN participants, we used an HCP-trained STIM to compute individual topological landscapes (see Methods). We initially examined the consistency of brain dynamics (low-dimensional embeddings after the filter function *f*) across development. The results showed a significant positive correlation between mean group consistency and age (Fig.5b), indicating that older individuals tend to achieve a more stable consensus in their brain dynamics. We also observed that age differences negatively correlate with global topology similarity between different age-sorted groups (see Methods), suggesting that groups closer in age exhibit higher similarities in their latent dynamics (Fig.5c).

We then investigated whether developmental differences were associated with specific movie content. Using the same STIM pipeline, we divided the time volumes into 41 non-overlapping cliques (mean duration = 14.6 seconds). Through manual labeling, we annotated the cliques with over 50% screen time featuring kids or minions (the small, yellow characters in the movie, as shown in Fig.5d). Subsequently, we generated the consensus of two extreme age groups (100 subjects for each group) and compared how individuals from intermediate ages deviated from this consensus as the age gap increased (the other 770 subjects). The lower part of Fig. 5e shows specific cliques where individuals deviated from the youngest group consensus (in orange) or engaged with the oldest group consensus (in blue) (*P* < 0.05 after FDR correction). Interestingly, we found that cliques causing individuals to deviate from the youngest consensus, compared to those engaging with the oldest consensus, featured significantly more scenes with kids or minions when calculated by time volumes (χ² (1) = 28.35, *P* < 0.0001, Fig. 5e), indicating that scenes rich in child characters are more likely to induce individual experiential differences among younger viewers during movie watching. Fig. 5f summarizes these observations, illustrating that younger individuals show greater heterogeneity and lower synchrony, with synchrony increasing progressively with age, especially in cliques featuring kids or minions. This indicates a marked development in global topology similarity.

We then examined the individual differences in global topology. First, we replicated the main results found in HCP data. A group-level shape graph was generated and annotated by the PC1 dimension on the global topological similarity matrix over time (top 5%: +VOLs; bottom 5%: -VOLs) (Fig.5g). Consistent with the HCP results, +VOLs exhibited significantly higher peripheral presence in the topological landscape (lower betweenness centrality: *t*=-4.61, *P*<0.0001) and an ‘attractor-like’ mode (higher populational synchrony: t=6.28, P<0.0001) compared to -VOLs. Given the dialogue-rich content of *Despicable Me*, we selected listening comprehension scores from the Wechsler Intelligence Scale (WIAT) as the cognitive metric. Across five equal-sized, non-overlapping age groups (Fig.5h), the principal gradient of global topology similarity significantly correlates with cognitive function in four groups (group-1: *r*=0.31, *P*<0.0001; group-2: *r*=0.08, *P*>0.05; group-3: *r*=0.15, *P*=0.032; group-4: *r*=0.36, *P*<0.0001; group-5: *r*=0.26, *P*=0.00031). In the HCP data, we observed differences between +VOLs and -VOLs in terms of movie content. However, due to the absence of natural scenes, we did not observe the significant opposite direction of cognitive correlation in HBN data.

For local geometry similarity, we focused on the paired correlations between cliques and mental health issues. From behavioral measurements provided by the HBN dataset, we selected seven cognitive dimensions and twelve psychological health dimensions with complete data, all adjusted to represent negative mental health outcomes (Supplementary Tab. 2). Using PLSR approach, we identified 14 probe-like cliques for behavioral correlation analysis. Consistent with the HCP findings, we discovered that specific movie content could explain the association between local geometry similarity and negative mental health (Fig. 5i). Typically, cliques associated with social mental problems primarily involve scenes of intense social interactions: three kids playing and yelling, refusing to go to bed (No.0, *ρ*=-0.15, *P*<0.0001), passing a gun to a bystander and then flying away (No.37, *ρ*=-0.12, *P*=0.0007), and a minion vocally protesting to Gru while attempting to escape (No.40, *ρ*=-0.13, *P*=0.0004). Anxiety-related issues appeared linked to scenes depicting negative parental reactions, such as a minion’s invitation being sternly rejected by Gru (No.34, *ρ*=-0.11, *P*=0.002). Emotional problems were potentially linked to extreme emotions: minions suddenly erupting in loud, prolonged laughter during play (No.18 *ρ*=-0.14, *P*=0.0002). Additionally, across all probe-like cliques, local geometry similarity was negatively correlated with negative mental health. The average correlation between +VOLs and negative mental health was significantly negative (*t*=-5.87, *P*<0.0001).

Given the rich social interactions in *Despicable Me*, we explored the potential application of local geometry in studying autism spectrum disorder (ASD). By conducting PCA on the behavioral measurements associated with ASD, we divided individuals into four groups based on progressively severe ASD-like symptoms excluding neurotypical controls (NC) individuals (n=165 for each group). We then compared the local geometry similarity of these groups with the NC group (n=63) across all probe-like topological cliques. For further insight, we also compared individuals officially diagnosed with ASD to NC individuals. We annotated the movie content and dialogue for the three cliques most significantly associated with ASD-specific behaviors. Interestingly, these cliques were all related to social behaviors (Fig. 5j). As ASD-like symptoms intensified, the difference in following behaviors compared to NC individuals progressively increased. This pattern suggests that customized movie scenes could potentially serve as neural ‘probes’ for identifying mental disorders such as ASD.

## Discussion

Naturalistic stimuli can elicit a continuous stream of rich subjective experiences through the adaptive coordination of neural systems. While watching a film is a collective experience(*10*, *13*, *16*), it also includes individual-specific elements likely determined by the viewer’s cognitive and psychological conditions(*34*, *35*). From a dynamical system perspective, we propose that individual divergence from group consensus is reflected in system-wide, low-dimensional brain latent dynamics, allowing for the probing of individual behavioral traits and mental disorders. To this end, we introduce a computational framework, STIM, designed to quantitatively elucidate the topological landscape of brain activity during naturalistic movie viewing. The STIM framework can generate a reliable group-level landscape of brain latent state dynamics and map individualized topological patterns into two distinct scales: global topology and local geometry, corresponding to individual divergence across and within specific movie cliques, respectively. Applying STIM to two large movie fMRI datasets, we demonstrated that the combination of rich naturalistic stimuli with brain recording holds great potential as neural ‘probes’ to measure individual traits in various behavioral domains. Specifically, global topology describes overall shape similarity and is specifically related to cognition: individuals who closely follow group dynamics in ‘attractor-like’ modes exhibit higher fluid intelligence, while those showing more diversity in ‘unconstrained’ modes also exhibit higher fluid intelligence. At the finer movie clique scale, local geometry explains multi-dimensional mental health traits beyond cognition. Lastly, we showed that STIM could capture the topological landscape related to developmental processes, with potential applications in detecting and understanding neurodevelopmental disorders such as autism.

STIM provides a novel method for analyzing brain states in the field of naturalistic fMRI, applicable to longer and more complex movie-viewing data. Compared to HMM, CAPs, and similar state-based methods, the TDA-based Mapper can represent a richer array of brain states during movie viewing. Additionally, it automatically and flexibly captures rapid transitions, preventing the blurring of distinct brain states, and provides user-friendly, dynamic visualizations. We observed that using the Mapper technique to construct low-dimensional topological landscapes independently in different subjects exhibits similar rapid transitions in response to the same movie content (Fig. 2b). STIM can further align these low-dimensional spaces across individuals to generate a group template, thereby mapping individual-specific states to behaviors.

Remarkably, this group-level topological landscape and the corresponding patterns of individual differences can be reliably established with a relatively small sample size (n=15). Thus, a cost-effective approach for future studies would involve collecting diverse movie clips from a small subset of participants to create a rich, group-level topological reference(*35*). The reference can then be used with plug-and-play movie segments tailored to various applications, guiding downstream scanning in larger populations. Although subjective evaluations during movie viewing are lacking, we speculate that STIM primarily captures the system-wide integration of brain activity during movie watching. This is especially evident considering that the synchronization of latent embeddings (using UMAP as the filter function) is higher than that original time series of visual regions, particularly in scenes requiring cognitive processing (Supplementary Fig. 2).

By applying the STIM framework to large-sample movie fMRI datasets, we have, for the first time, elucidated the individual differences in latent dynamics during watching rich movie contents. In terms of global shape-like topology, our findings reveal that different naturalistic stimuli elicit either common or heterogeneous group consensus. Furthermore, individual divergence is specifically correlated with cognitive abilities, but the direction of this association is context-dependent. The principal divergence in global topology spans multi-level domains, including topological attributes within the brain’s landscape (e.g., shape graph), synchronous modes within the population (e.g., interindividual distance), brain configurations (e.g., spatial networks), movie contents (e.g., stimuli categories), and individual traits (e.g., behavioral measures). To enhance interpretability, we decomposed the first component of the global topology metric into two distinct temporal parts: +VOLs, representing top positive contributions, and -VOLs, representing top negative contributions. +VOLs correspond to more complex scenes, including characters and dialogues, aligning with our initial hypothesis of stimuli as anchors or attractor basins guiding shared low-dimensional latent dynamics across individuals. These attractors are distributed in the brain’s landscape periphery and correspond to multiple network-dominant brain configurations. During movie-watching, individuals whose landscapes can flexibly ‘follow’ these +VOLs attractor-like states exhibit higher fluid intelligence. Based on PCA approach, Shine et al demonstrated a common attractor-like space facilitating the execution of multiple cognitive tasks, with individuals having greater fluid intelligence tending to exhibit more effective flow through the space(*19*). We speculate that +VOLs may be homologous to this attractor-like space from a dynamical system perspective, but corresponding to more multifaceted, naturalistic cognitive processes. In contrast, -VOLs, which correspond to more natural scenes, occupy the brain’s landscape center and exhibit more mixed network configurations. Seemingly similar patterns have been previously noted in the analysis of dynamic brain states. For instance, Saggar et al. used a TDA-based Mapper to reveal the topological center in individual landscapes during the resting state, suggesting that the uniform state acts as an intermediary to modulate transitions between other network-dominant states(*25*). However, differed from previous studies, the -VOLs characterize the overall patterns across the group, thus averaging out the heterogeneous brain activations of many individuals. At the individual level, -VOLs could still encompass numerous network-dominant volumes, but statistically, the mixed network level is significantly smaller than that of +VOLs. Interestingly, we have discovered for the first time that under specific stimuli (-VOLs), individuals with more divergent traversals from group-level latent dynamics exhibit higher fluid intelligence. This finding suggests the importance of spontaneous divergent thinking, similar to mind wandering, especially when encountering stimuli that lack explicit direction and cognitive load. Considering that the HBN dataset lacks natural-themed content, the first component of the global topology metric consistently has positive weights and can reliably explain individual differences in cognitive dimensions across different age groups.

A conceptual prospect of the naturalistic paradigm is to use designed movie fragments to induce rich brain activation, with the expectation that different fragments can assess various dimensions of behavior or disease symptoms. This potential has been suggested by previous studies, which showed that certain movie stimuli (e.g., independent video clips and the presence of faces) can predict sex(*36*), cognition(*7*), or emotion domains(*37*) to varying degrees. At a finer scale, our results suggest that specific movie scenes can serve as ‘probes’ to detect corresponding behaviors. This is facilitated by the STIM framework, which can segment complex movie content into local geometry clusters based on brain latent dynamics. Importantly, our reported clique-behavior pairs are largely explainable. For instance, the cliques most correlated with life satisfaction predominantly feature pleasant and peaceful conversations, while the two scenes associated with anger traits both contain unfriendly language, such as a note with ‘udick’ and a conversation with the word ‘damn’. In the HBN dataset, the contagious laughter of the Minions clique corresponds to individual differences in the Affective Reactivity Index, a psychological assessment used to measure emotional reactivity. The clique-behavior explainability highlights that movie scenes can be further strategically designed to target specific dimensions of mental health(*35*, *38*), reducing the subjectivity of self-report measures(*7*, *39*). In most cliques, negative mental health issues disrupt individuals from engaging corresponding group-level latent dynamics; within HCP -VOLs, scenes of natural, eerie forests show that individuals with greater divergence from the group tend to have a lower tendency towards negative traits (i.e., loneliness).

Our results suggest the potential of the STIM framework to detect developmental issues or psychiatric disorders and inspire intervention strategies. For the same movie stimuli, groups of similar ages exhibit comparable global topology similarity and become more synchronized within the group as they develop. Different developmental groups show varying latent dynamics, such as cliques relating to different characters. Individuals with autism significantly deviate from healthy controls in latent dynamics, especially in social scenes. Previous works have suggested that the brain activity of individuals with psychiatric disorders during naturalistic stimuli shows abnormalities compared to healthy controls(*40–42*). In the future, constructing landscape references of healthy groups from different age brackets to naturalistic stimuli as a normative model may enable real-time observation of individual divergence from demographic-specific templates using the Mapper visualization advantage. For example, an individual at high risk for autism may specifically deviate in certain social scenes, such as interactions with parents rather than peers. Understanding these deviations could potentially help customize behavioral intervention strategies at the individual level.

We consider several limitations of the current work. First, STIM framework can be improved in several ways. For filter function, we used UMAP for its ability to preserve topological structure of original data, but its representational capacity is limited. Trainable deep learning models, such as the recently proposed CEBRA approach(*43*), could produce higher performance latent spaces. For binning and clustering, we used the traditional Mapper approach. However, newer methods like NeuMapper suggest that the binning and clustering processes can be modified to align with the temporal dynamics of fMRI data(*44*). Second, our study is based on whole-brain regions, but for specific subdomains like the emotion domain, selecting key nodes in emotional circuits as inputs may yield better behavioral correlations(*45*, *46*). Third, the STIM framework inherently studies brain activity in low-dimensional representations, which cannot resolve cross-region synchrony. This can be addressed using classical ISC methods. Fourth, the HBN dataset contains a very small proportion of healthy samples and has a wide age distribution, limiting the construction of models for classifying healthy controls and patients. Future studies could build age-matched disease cohorts with movie fMRI to explore the potential for individual-level diagnosis. Fifth, future studies could employ more convenient data collection methods, such as electroencephalograph (EEG) or magnetoencephalography (MEG).

## Method

### Dataset 1: Human Connectome Project (HCP) dataset Participants

The movie-watching fMRI (mv-fMRI) data of healthy young subjects was obtained from the HCP database(*47*, *48*). We analyzed 170 healthy participants who completed the movie-watching sessions (average age = 29.4 years, SD = 3.3 years; 105 females). Notably, the dataset contained twins and siblings, hence participants were from 89 unique families. In our validation tests, we mitigated potential biases from familial relationships. Ethics approval was obtained by the authors of the original studies in accordance with the Washington University Institutional Review Board (IRB), and the Washington University–University of Minnesota (WU-Minn HCP) Consortium ensured full informed consent from all participants.

### Movie information

For details of the HCP movie protocol, please refer to the Human Connectome Project protocols for 7T imaging (https://www.humanconnectome.org/hcp-protocols-ya-7t-imaging). Briefly, participants viewed audiovisual movie clips in 4 runs across two separate sessions, with each run consisting of 4-5 distinct movie segments. There was a 20-second rest period between clips during which the word “REST” appeared on the screen. Audio was delivered through Sensimetrics earbuds. In our analysis, we excluded movie segments shorter than 2 minutes, resulting in 13 clips from various sources including independent films and Hollywood movies, providing a rich array of naturalistic stimuli. These segments covered multiple genres such as sci-fi (*Inception*), drama (*Two Men*; *Social Network*; *Ocean’s Eleven*; *Home Alone*; *Erin Brockovich*; *Empire Strikes Back*), romance (*1212*), documentary (*Welcome To Bridgeville; Mrs Meyers Clean Day*; *Pockets*), music video (*Off The Shelf*) and nature scenes (*Northwest Passage*) (for detailed information on the movie segments, see Supplementary Tab. 1). To correlate with the real-time viewing experience, we used manually assigned semantic labels provided by HCP (7T_movie_resources/WordNet_MOVIE1_CC1.txt, etc.). These labels identified objects (e.g., road, men, tree) and human actions (e.g., talk, sit, walk) depicted in the movie scenes, with label frequency aligned to the volumes (i.e., providing an array of labels every second).

### Data acquisition

All movie-watching fMRI data were acquired using a 7 Tesla Siemens Magnetom scanner with a 32-channel head coil. Among the mv-fMRI scans conducted across four runs, runs 1 and 4 were acquired with anterior-to-posterior (AP) phase encoding, while the remaining two runs used posterior-to-anterior (PA) phase encoding. All mv-fMRI scans were collected using a gradient-echo-planar imaging (EPI) sequence with the following uniform parameters: repetition time (TR) = 1000 ms, echo time (TE) = 22.2 ms, flip angle = 45 degrees, field of view (FOV) = 208 × 208 mm (RO x PE), matrix size = 130 × 130 (RO x PE), slice thickness = 1.6 mm, resulting in 1.6 mm isotropic voxel resolution. Additional scanning parameters included a multiband factor of 5, an image acceleration factor (iPAT) of 2, partial Fourier sampling of 7/8, echo spacing of 0.64 ms, and a bandwidth of 1924 Hz/Px. Structural T1-weights (T1w) scans were acquired using a 3T Siemens Connectome Skyra scanner, with the following parameters: TR=2400ms, TE=2.14ms, TI=1000ms, flip angle=8 degrees, FOV=224 × 224 mm, voxel size=0.7mm isotropic, iPAT=2, bandwidth=210 Hz/Px.

### fMRI preprocessing

We initiated our analysis using minimally preprocessed data (*Movie Task fMRI 1.6mm Functional Preprocessed*)(*49*). The HCP minimal processing is a standardized preprocessing pipeline tailored for the HCP dataset designed to minimize information loss. This includes gradient distortion correction, motion correction, fieldmap-based EPI distortion correction, EPI to T1w registration, non-linear registration (FNIRT) into MNI152 space, and intensity normalization. Additionally, we performed nuisance signal correction by regressing out Friston-24 motion parameters(*50*), 3 temporal trending parameters (constant, linear, and quadratic trends), as well as signals from white matter (*WMReg.2.nii.gz*), cerebrospinal fluid (*CSFReg.2.nii.gz*), and the average global signal extracted from the brain mask (*brainmask_fs.1.60.nii.gz*). Lastly, a temporal band-pass filter (0.01Hz<f<0.08 Hz) was applied to the data.

### Dataset 2: Healthy Brain Network (HBN) dataset

#### Participants

We utilized the HBN dataset, a large-scale multimodal dataset focusing on the development and mental health of children and adolescents, collecting neuroimaging and behavioral data from various age groups in the New York area(*51*). Data selection was based on the completeness of imaging and behavioral data, correctness of preprocessing, anomalies in head motion and synchrony (see below). This resulted in a subset of 970 participants (ages 5-21, average age = 10.6 years, SD = 3.5 years; 341 females). Among these, 95 individuals had no diagnosis given named as normal control (NC), while the majority were diagnosed with one or more psychological issues. The Child Mind Institute confirmed that adult participants provided full informed consent, and minors aged 5-17 gave assent, with their parents completing the full informed consent. For more details, please refer to: https://fcon_1000.projects.nitrc.org/indi/cmi_healthy_brain_network/index.html.

### Movie information

The HBN dataset provides fMRI scans from two different movie segments watched with simultaneous audio: *The Present* (3 minutes 21 seconds) and a clip from *Despicable Me* (10 minutes). ‘The Present’ narrates the story of a boy with an amputation who receives a three-legged puppy from his mom and eventually warms up to it. The *Despicable Me* clip recounts various events involving three children and the supervillain Gru, including Gru tucking the children into bed, discussing plans to steal the moon with Dr. Nefario, the children being returned to the orphanage, and Gru rocketing into space. Given that *The Present* is shorter and lacks dialogue and plot transitions, we chose to analyze only the data from the *Despicable Me* clip.

### Data acquisition

Structural and functional MRI data were collected from two different sites: the CitiGroup Cornell Brain Imaging Center (CBIC) using a Siemens 3T Prisma scanner and the Rutgers University Brain Imaging Center (RUBIC) using a Siemens 3T Tim Trio scanner, both employing a 64-channel head coil with identical scanning parameters. Mv-fMRI scans were conducted with the following parameters: TR=800ms, TE=30ms, flip angle=31 degrees, resolution=2.4 mm × 2.4 mm × 2.4 mm, 60 slices, FOV phase=100%, and a multi-band factor of 6. T1w scans were collected with: TR=2500ms, TE=3.15ms, TI=1060ms, flip angle=8 degrees, resolution=0.8 mm × 0.8 mm × 0.8 mm, 224 slices, and FOV phase=100%. We did not perform inter-site corrections or analyses during data processing; instead, site information was treated as a covariate and regressed out in subsequent behavioral analyses.

### fMRI preprocessing

Anatomical and functional data preprocessing was conducted using fMRIPrep 20.2.1(*52*). The T1w image underwent intensity non-uniformity (INU) correction with N4BiasFieldCorrection, provided by ANTs 2.3.3, and it was designated as the reference image. This reference T1w image was then skull-stripped using a Nipype implementation of the antsBrainExtraction.sh script from ANTs, utilizing the OASIS30ANTs template for anatomical accuracy. Following this, segmentation of brain tissues—specifically CSF, WM and GM—was performed on the skull-stripped T1w image using the FAST tool from FSL 5.0.9. The volume was spatially normalized to the MNI152NLin2009cAsym standard space through nonlinear registration using antsRegistration from ANTs 2.3.3, ensuring consistent anatomical alignment across studies.

For each subject, a reference volume and its skull-stripped version were generated. The BOLD reference was aligned (co-registered) to the anatomical T1w reference using the FLIRT tool from FSL 5.0.9. Motion correction was applied, and the data were resampled and normalized to the standard space (MNI152NLin2009cAsym). Metrics such as Framewise Displacement (FD), CSF, WM, and global signal were calculated. Noise reduction on the BOLD signals was performed using the CompCor method.

Similar to the preprocessing of HCP, we performed nuisance signal correction by regressing out Friston-24 motion parameters, three temporal parameters (constant, linear, and quadratic trends), as well as signals from WM, CSF, and the global signal. Lastly, a temporal band-pass filter (0.008Hz<f<0.15 Hz) was applied to the data. After data processing, 1,227 subjects had complete imaging data (or were correctly preprocessed) along with basic information (age, gender, disease diagnosis). Considering the propensity for greater head movement in children and adolescents, we excluded 159 individuals with excessive head motion (mean FD > 1mm). We also examined the average head motion in each fMRI run from the HCP database (Movement_RelativeRMS_mean.txt). We did not exclude any subjects from the HCP dataset because none of the runs showed a mean FD exceeding 0.5 mm. We also removed 98 individuals whose data exhibited abnormal synchrony (average Pearson correlation with the group mean < 0.1 across 271 ROIs, whereas the lowest in HCP is 0.12). We suspect these individuals were not engaged in the movie-watching task (Supplementary Fig. 7)

### Extraction of whole-brain time series

Consistent with our prior work(*53*), we selected 271 predefined regions of interest (ROIs) spanning the entire brain. We obtained 200 cortical regions from the Schaefer atlas(*54*), 54 subcortical regions from the Melbourne atlas 7T version scale III (3T version scale IV for HBN)(*55*), and 17 cerebellar networks from the Buckner atlas(*56*). Among the cortical ROIs, seven different networks were assigned based on previous work(*54*), including the visual network (for short: Visual), somatomotor network (SomMot), dorsal attention network (DorsAttn), ventral attention network (SalVentAttn), limbic network (Limbic), frontoparietal control network (Control), and default mode network (Default). The subcortical and cerebellar ROIs were treated as a whole, without further network subdivision. Considering the hemodynamic response delay, we shifted all time-series signals from movie-viewing by 5 seconds to correspond to 5 TRs in HCP(*57*). As the HBN movie-viewing protocol does not include resting periods before and after the movie, and thus does not allow for temporal shifts in the time series, we compensated by delaying the alignment of movie-related labels with the BOLD time series by 5 seconds (equivalent to 6 TRs).

### Classic Mapper

We utilized the Mapper pipeline developed by Veen et al(*58*) and Geniesse et al(*26*). to generate the shape graphs. The classic Mapper pipeline includes the following steps: i) *Dimensionality reduction (Filtering)*: The high-dimensional data are embedded into a lower dimension (d=3) using a filter function. We chose UMAP (*n neighbors* = 25) to preserve the topological features of the data, utilizing PCA for initialization to maintain global structural integrity. ii) *Overlapping binning*: This stage involves overlapping three-dimensional binning to encapsulate data points, with a resolution parameter (#bins) set to 12, resulting in a total of 12 × 12 × 12 = 1728 *bins*, and a 50% overlap between bins. iii) *Partial clustering*: Within each bin, partial clustering groups data points into nodes based on the original high-dimensional information. We utilized density-based spatial clustering of applications with noise (DBSCAN) (*59*), which does not require a predetermined number of clusters. 4) *Shape graph generation*: The Mapper approach treats each cluster within the bins as a node (containing multiple volumes). Overlapping bins cause some nodes to contain identical volumes, thereby generating edges between nodes.

Taking individual shape graphs shown in Fig. 1b and 2b as example, the input data consisted of parcellated time series for each participant; for Fig. 1b, we used 7 movie clips with dimensions of 1439 *volumes* × 271 *ROIs*, and for Fig. 2b, the *Inception* clip with dimensions of 226 *volumes* × 271 *ROIs*. The Mapper pipeline outputs a graph object. The shape graphs can be annotated with various meta-information, such as nodal degrees, network configuration, and other relevant metrics, enabling a comprehensive understanding of the brain latent dynamics.

### STIM framework

Building on the classic Mapper, we introduce the STIM framework to analyze individual differences in brain latent dynamics during naturalistic movie watching. The STIM framework posits a group consensus of latent dynamics, akin to the ‘Anna Karenina model’ (derived from Tolstoy’s opening line: “All happy families are alike; each unhappy family is unhappy in its own way”), and quantifies each individual’s dynamic trajectory deviations from the group consensus(*13*). Practically, STIM integrates whole-brain spatiotemporal activity from multiple participants, unfolding and aligning individual latent dynamics to establish a group-level reference as a proxy of consensus. By comparing the global and local topological differences between individual subjects and the consensus, STIM provides two distinct metrics of individual differences: global topology and local geometry similarities.

We applied a joint dimensionality reduction approach to achieve group-level low-dimensional dynamics. For HCP data, we stacked the whole-brain time series data of all individuals (dimensions: from 170 *subjects* × 2804 *volumes* × 271 *ROIs* to 476680 × 271 *ROIs*) and performed dimensionality reduction along the ROI axis, thereby creating an aligned low-dimensional space. From this, we extracted each individual’s low-dimensional embedding (dimensions: 2804 *volumes* × 3) and standardized the embedding by z-score normalization across volumes. The low-dimensional embeddings were then averaged to the group-level dynamics of consensus. We tested three common dimensionality reduction methods (or named filter *f* functions): PCA, t-SNE, and UMAP. By comparing the consistency of group-level dynamics across different participants, we finally selected UMAP as the filter function *f* (Supplementary Fig. 8). In the STIM framework, we then integrate individual and group-level low-dimensional dynamics (dimensions: (2804 + 2804) *volumes* × 3), to proceed with the TDA-based Mapper steps, which include *Overlapping binning*,

*Partial clustering*, and *Shape graph generation*. Considering the HCP movie stimuli contain richer themes, we incorporated HCP data to generate 3-dimensional UMAP trajectories when constructing low-dimensional representations for HBN.

### Temporal connectivity matrix

As described above, we aligned the complex latent dynamics of different participants within a shared topological space, generating a comprehensive Mapper shape graph. Each node in the shape graph contains a set of time points considered similar. Following the previous work(*24*), we quantified the topological similarity between time points by utilizing the clustering relationships within nodes and the connections between nodes, resulting in a Temporal Connectivity Matrix (TCM, denoted by *T*). Element *T*_*ij*_ within *T* quantifies the connectivity strength between time frames *VOL*_*i*_ and *VOL*_*j*_.

The connectivity strength, *T*_*ij*_, is calculated as the ratio of the cardinality of the intersection set of nodes shared between time frames *VOL*_*i*_ and *VOL*_*j*_ relative to the maximum cardinality of the intersection of nodes involving *VOL*_*i*_ and any other time frame *VOL*_*p*_ across the entire dataset. Symbolically, this is expressed as:

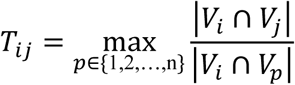

where ‘|·|’ indicates the cardinality of a set, *V*_*i*_ and V_j_ are the sets of nodes involving with time frames *VOL*_*i*_ and *VOL*_*j*_, respectively, and the denominator is the maximum cardinality obtained by the unite of *V*_*i*_ with every other set *V*_*p*_ across all time points *p*.

The resulting matrix *T* is symmetrized by:

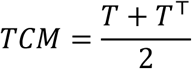

This matrix provides a measure of topological similarity on a fine temporal scale. Notably, topological similarity is a discrete measure, therefore, we use Spearman rank correlation for subsequent association analyses.

In the classic Mapper, the TCM computed for a single subject is dimensioned as *n volumes* × *n volumes*. In STIM, which integrates group dynamics, the resulting TCM dimensions expand to 2*n volumes* × 2*n volumes*. From this, we extract the intersecting matrix to measure topological similarity between different individuals (Supplementary Fig. 9).

### Individual divergence

Features were extracted from the TCM across multiple scales. Taking HCP as an example, at the global scale, we extracted the diagonal of the TCM, defined as global topology similarity (dimensions: 170 *subjects* × 2804 *volumes*). This metric quantifies the overall shape differences between individual and group trajectories. In the STIM framework, due to discontinuous overlapping bins, global topology similarity is discrete value. This discrete, or ‘fuzzy’, metric is potentially appropriate for the analysis of biological signals, which are inherently imprecise. Within the context of movie viewing, we consider global topology similarity could reflect the extent to which an individual’s subjective experience aligns with the group consensus.

At the local scale, the TCM matrix visually exhibits a modular structure, suggesting that this low-dimensional representation can serve as a basis for temporal segmentation. A standard change point detection algorithm, pruned exact linear time (PELT)(*31*), was employed to segment the TCM matrix based on group-averaged dynamics. Notably, we excluded the 50% subjects whose data demonstrated lower similarity during the averaging process, to ensure that the segments were not influenced by outliers. The parameters for PELT were selected using a data-driven approach, with the evaluation metric being the degree of clustering modularity, as proposed by Busch, E.L., et al(*60*). Specifically, we chose the PELT parameters that resulted in the maximum ‘within-versus between-event difference score’, or in other words, the highest boundary sharpness. Using PELT algorithm, we identified 161 non-overlapped cliques in the HCP data (mean duration=16.2s, SD=9.0) and 41 non-overlapped cliques in the HBN data (mean duration=13.7s, SD=7.0). Clique segmentation information was mapped back onto individual TCMs, from which submatrices corresponding to distinct cliques were extracted. These submatrices were then combined with the group average clique through a weighted summation to compute the local geometry similarity for individuals within specific cliques. Local geometry similarity reflects an individual’s alignment with the group consensus for specific movie content. Compared to the sliding window approach, this method avoids averaging distinctly different time points together.

HCP provided a manually labeled annotations for each second, which can be interpreted as subjective understanding for movie scenes. For example, a classic annotation can be [‘man’, ‘run’, ‘road’, ‘sky’]. Utilizing these semantic labels, we calculated a semantic temporal connectivity matrix (semantic-TCM), where the elements *T*_*ij*_ represent the proportion of overlapping word entries between semantic vectors *T*_*i*_ and *T*_*j*_. The semantic-TCM was then input into the PELT algorithm to segment it into non-overlapping semantic cliques. Subsequently, we computed the minimized linear error between segments of topological cliques and segments of semantic cliques using a standard algorithm for solving the linear assignment problem(*61*). For example, the minimized error between segment [5,14,21] with segment [4,15,20] is 3. For statistical testing, we generated a null model by randomly segmenting the temporal axis with the same number of segments and repeating this process 10,000 times.

### Robustness of STIM framework

We performed two additional analyses to ensure the robustness of the STIM framework:

1. Robustness of group-level dynamics: We randomly sampled two independent groups of participants and comparing their averaged group-level dynamics. To enable a fair comparison, time series data were also derived from three randomly selected ROIs. The individuals in two sampled groups had no familial relationships. We generated UMAP low-dimensional embeddings of group references ranging from 5 to 40 individuals and found that the group reference consistency, measured by Pearson correlation, was significantly higher than that of the original time series (Fig. 2e). Moreover, we found that for most movies (8 out of 13), the group consistency of whole-brain latent dynamics was significantly higher than that achieved using even the top three synchronized ROIs in the visual network (Supplementary Fig. 2).
2. Robustness of individual difference: we calculated the global topology using group-level dynamics based on smaller groups of subjects and compared it to that using all subjects. We found that with a smaller sample size ( *n subjects* = 15), the Pearson correlation of individual differences to the group reference formed from all subjects (*n subjects* = 170) averaged 0.98 across multiple samplings (Fig. 2f).

### Parameter selection and perturbation analysis

Similar to the method used by Saggar et al.(*24*) to determine the Mapper parameters, we optimize the resolution parameter by maximizing the topological individual differences. Specifically, we computed the standard deviation of global topology and local geometry across individuals, to reflect how different resolution parameter span the individual differences effectively. Ranging from fuzzy resolution (*n bins* = 4) to fine-grained resolution (*n bins* = 20), we select the resolution parameter *n bins* = 12 for maintaining top standard deviation and high resolution. Supplementary Tab. 5 shows the standard deviation across different resolution parameters.

Previous work(*24*) has demonstrated the robustness of the Mapper approach to parameter perturbations. Once again, we tested the impact of a wide range of Mapper parameters on the main outcomes of individual differences using HCP data. We varied the resolution parameter, calculating the Pearson correlation of global topology (as well as local geometry) between perturbated resolution and selected resolution (*n bins* = 12). The results indicate that under most parameter settings, the topological properties and the outcomes related to individual differences remain stable (Supplementary Fig. 10).

Further, utilizing the criteria proposed by Hasegan et al.(*62*), we examined the validation of Mapper shape graph. Briefly, a valid Mapper shape graph needs: 1) cover most of the input data (coverage beta *β* > 70%); 2) captures more than trivial autocorrelation dynamics (non-autocorrelated nodes *α* > 15%, with autocorrelation threshold *τ* = 11); 3) has a non-trivial structure (pairwise distances entropy *S* > 2). Here, we computed these three criteria in HCP group shape graph (*β* = 99.64%, *α* = 87.61%, *S* = 2.12), ensuring the effectiveness of the generated shape graph.

### Behavioral items selection

We hypothesize that the global geometric similarity corresponding to topological cliques is related to participants’ cognitive or comprehension abilities, while local geometric similarity can explain multi-dimensional human behaviors in a manner related to specific movie content. To test this hypothesis, we utilized behavioral questionnaires from the HCP and the HBN datasets.

*HCP Dataset.* In the HCP dataset, we included 59 behavioral measurements spanning multiple domains, including cognition, emotion, psychiatric, and life Function. The specific domains incorporated into our analysis are detailed in Supplementary Tab. 2. Given our hypothesis that understanding movie content primarily relies on abilities such as problem-solving, logical reasoning, and pattern recognition, we chose Fluid Intelligence as the key cognitive measure for our study, rather than crystallized intelligence or executive function. Fluid Intelligence was assessed using a shortened version of Raven’s Progressive Matrices, specifically the A table, developed by Gur and colleagues(*63*).

For subjective emotional and psychological health measurements, we selected self-report questionnaires from the Emotion domain and Life Function (Achenbach Adult Self-Report, Syndrome Scales, and DSM-Oriented Scale). We did not include task-based measurements (for example emotion recognition) or psychiatric history because self-report questionnaires and life function metrics more subjective, matching our ‘consensus and divergence’ hypothesis. For clarity, we inverted the scores for positive items in emotional and psychological tests, including life satisfaction (*LifeSatisf_Unadj*), meaning and purpose (*MeanPurp_Unadj*), positive affect (*PosAffect_Unadj*), friendship (*Friendship_Unadj*), emotional support (*EmotSupp_Unadj*), instrumental support (*InstruSupp_Unadj*), and self-efficacy (*SelfEff_Unadj)*, ensuring consistency in the direction of negative emotions and psychological health expressions.

*HBN Dataset.* In the HBN data, we firstly aimed to replicate the assessment of cognition and mental health issues similar to the HCP. Beyond HCP, the HBN recruited children and adolescents aged 5-21 and provided clinical diagnostic labels for mental disorders, this allowed us to further explore specific issues related to adolescent mental health development and disorders. HBN participants exhibited a variety of transdiagnostic conditions (e.g., autism spectrum disorder), providing a comprehensive context for studying developmental trajectories and the spectra of mental disorders.

Specifically, in the cognitive dimension, given the lack of a direct measure of fluid intelligence in the HBN and the dialogue-based nature of movie content, we chose the listening comprehension subtest (WIAT_LC_Stnd) to evaluate cognitive comprehension abilities. For results of other cognitive behavioral tests provided in the HBN dataset, please refer to the Supplementary Tab. 3.

For mental health, self-report scales were used to assess subjective emotional experiences and mental health status, with data adjusted to reflect negative outcomes. We focused on participants older than 8 years, due to the instability of mental health issues in very young subjects(*51*), and excluded questionnaires with fewer than 500 responses. This resulted in the identification of 12 mental health-related items (Supplementary Tab. 2).

For the analysis of autism spectrum disorder (ASD), we first classified each participant as either ‘Disorders’ (n=662) or ‘No Diagnosis’ (n=63) based on the clinical diagnostic information from the HBN sample. We then identified the ASD group from the ‘Disorders’ subjects in two ways: 1) groups with increasingly severe ASD-like symptoms: We computed an indicator to estimate the severity of ASD-like symptoms.

Specifically, we conducted a principal component analysis on behavioral measures related to ASD, including the Autism Spectrum Screening Questionnaire, Social Communication Questionnaire, Social Responsiveness Scale, and Repetitive Behavior Scale. We used the first principal component (explained 71.6% of the total variance) as the main indicator of ASD-like symptoms and divided individuals into four equal-sized subgroups based on the severity of these symptoms (n=165 per group). 2) Autism diagnosis group: We used diagnostic labels to classify subjects into the Autism diagnosis group. Notably, the Autism diagnosis group is a subset of the ASD-like symptoms group. By comparing differences between the Autism diagnosis group and the ASD-like symptoms group with the ‘No Diagnosis’ group, we aimed to provide insights into the potential application of biomarkers in the development process of Autism.

### Individual difference in global topology

We conducted PCA on the global topology similarity of individuals (HCP data: 170 *subjects* × 2804 *volumes*) along temporal volumes. The top 5% positive volumes are referred to as +VOLs, and the bottom 5% negative volumes as -VOLs. In the HBN data, we divided participants into five equally sized age groups (the age range of five groups are: [5-8, 8-9, 9-11, 11-14, 14-21], dimensions : 5 × 194 *subjects* × 750 *volumes*) and then conducted PCA to calculate the PC1 for each group separately, avoiding the effects resulting from systemic changes during development.

### Betweenness centrality

We applied the betweenness centrality from graph theory to estimate the topological attributes of the shape graph. Betweenness centrality measures a node’s role in facilitating information flow within the network by quantifying its presence on the shortest paths between other nodes(*32*).

Mathematically, the betweenness centrality *C*_*B*_(*v*) for a node *v* is defined as:

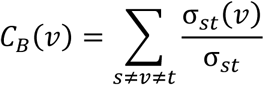

where σ_*st*_ is the total number of shortest paths from node *s* to node *t* and σ_*st*_(*v*) represents the number of those paths that pass through *v*. This measure sums the fraction of all shortest paths that pass through a given node across all pairs of nodes. Due to the potential overlap of volumes across nodes in the shape graph, we calculated the average betweenness centrality of these nodes to determine their corresponding volumes’ centrality.

### Group synchronous modes

We averaged the global topology similarity across all individuals to identify the population synchrony at different volumes. The synchrony in +VOLs was significantly higher than in -VOLs, indicating two distinct synchronous modes. Additionally, in the HCP data, we observed a linear positive correlation between volumes population synchrony and the weights of PC1 (Pearson *r* = 0.48, *P* < 0.0001). The consistency between individual difference gradient (PC1 of global topology) and synchronous modes indicates that individuals who ‘follow’ the group consensus tend to do so consistently across different volumes.

### Network configuration

Adapting methods from Saggar et al. on resting-state networks, we examined network configurations in mv-fMRI(*25*). The network configuration is determined by the overall activation pattern within a shape graph node, visualized in Mapper as the proportion of a pie chart for each node. Specifically, we normalized whole-brain activities in each volume, and calculated the values for nine networks (Yeo 7 network(*54*), subcortical region(*55*), and cerebellum(*56*)). To determine whether a dominant network configuration exists within nodes, we calculated a node’s network mixed index as 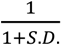, where *S. D*. represents the standard deviation of the activation levels across different networks. A higher network mixed index signifies more uniform activation across networks. In the visualization of the Mapper shape graph (Fig. 3d, e), the proportions in the pie chart for the network only include the positive values after normalization.

### Network contribution to global individual differences

The contribution of each ROI to global topology similarity was separately analyzed for +VOLs and -VOLs. Taking +VOLs as an example, we first concatenated the time series data of +VOLs (the dimension: 170 *subjects* × 271 *ROIs* × 140 *volumes*). For each ROI, we computed the Pearson correlation between the subject and group averaged time series (denoted as ROI similarity, the dimension: 170 *subjects* × 271 *ROIs*). For each ROI, we calculated the Pearson correlation between ROI similarity (the dimension: 170 *subjects* × 1) and the averaged global topology similarity of +VOLs (the dimension: 170 *subjects* × 1). This measure represents the ROI’s contribution to the individual divergence observed in +VOLs.

### Selection of probe-like cliques

Considering the high-dimensional nature of both cliques and behavioral dimensions, we adopted a two-stage approach to identify statistically significant pairs: 1) Identification of statistically significant cliques with behavior: We performed PCA on 59 behavioral items, using top 5 PCs that explained 49.6% of the total variance in HCP data (71% in HBN data). Subsequently, we used Partial Least Squares Regression (PLSR) to estimate the linear relationship between individual local geometry similarities (dimension: 170 *subjects* × 1) and behavioral PCs (dimension: 170 *subjects* × 5). We then selected cliques that demonstrated statistical significance in permutation tests (P < 0.05, n = 10,000 permutations), with each permutation involving a shuffle of subject IDs across local geometry similarities. 2) Behavioral correlation analysis: we calculated the Spearman correlation between individual local topological similarities and single behavioral items, controlling for age and gender as covariates. This step measures how specific cliques explain specific behaviors.

### Cliques-Mental health pair wise analysis

After identifying probe-like cliques and their pairwise relationships with specific behaviors, we analyzed the behavioral dimensions of mental health. We assessed the ‘gradient’ under the mental health dimension by averaging the significance of probe-like cliques. The derived weights for different behaviors were visualized using a word cloud (Fig.4f).

In presentation focusing on clique-mental health pairs, we first concentrated on the two behaviors with the highest weights—*loneliness* and *life dissatisfaction.* We highlighted the four cliques most significantly associated with each of these behaviors. The displayed P-values are the raw Spearman correlation P-values between each behavior and the local geometry similarity of the respective cliques. Among all probe-like cliques, we displayed the other representative clique-behavior pairs; for a complete list, please refer to Supplementary Tab. 4. Notably, we also displayed some significant clique-behavior pairs that did not pass the permutation test, including No.57, No.108, No.78, and No.128.

### Dual directions of dynamics modes

To further investigate how negative mental health affects individual divergence, we examined if the correlation direction differed between ‘attractor-like’ modes and ‘unconstrained’ modes. We extracted the global topology similarity (170 *subjects* × 2804 *volumes*) in +VOLs and -VOLs (n volumes = 140), resulting in ‘attractor-like’ and ‘unconstrained’ modes for individual topological scores (2 × 170 *subjects* × 140 *volumes*). We then selected the top 10 negative mental health behavioral metrics (170 *subjects* × 10 *behaviors*), based on the weights shown in the word cloud (Fig.4f), including life satisfaction, loneliness, ASR somatic problems, ASR internalizing, perceived rejection, DSM somatization, sadness, perceived stress, DSM anxiety, and anger hostility. Subsequently, we computed the Spearman correlation between global topology similarity and negative behaviors (for +VOLs or -VOLs, the dimension is 140 *volumes* × 10 *behaviors*). Finally, we performed a one-sample t-test with a mean of zero on the average Spearman correlations for +VOLs and - VOLs (dimension: 140 *volumes* × 1, Fig. 4h).

### Developmental analysis

In the development-related analysis (Fig. 5b-h), we utilized the shared low-dimensional space (UMAP) trained from HCP data and independently transformed individuals from the HBN data. We opted not to apply dimensionality reduction directly on the HBN dataset due to the following considerations: (i) The age distribution is highly skewed, particularly with a concentration of participants between the ages of 8 and 10, which could introduce bias into the shared topological space. (ii) Distinct latent dynamic patterns may emerge at different developmental stages, and applying joint dimensionality reduction could obscure the unique characteristics specific to each age group.

To assess the consistency of low-dimensional dynamics across developmental stages, we used one-year intervals as the unit of analysis. For each interval, we calculated the Spearman correlation of low-dimensional embeddings between two independent, non-overlapping samples (15 subjects per group). Fig.5b displays the mean values along with the 95% confidence intervals for the consistency, based on 1000 iterations of random sampling within each interval. Further, we explored developmental patterns of different group consensus (Fig.5c). Considering the uneven distribution of participants across different age groups, we grouped participants into 10 non-overlapping groups of equal size (10 × 97*subjects*) to ensure a balanced analysis. For consensus analysis, we first calculated the consensus within each group using the STIM framework. Then, we computed and averaged the global topology similarity for each pair of groups. Age difference indicates the intervals between two groups, for example the age difference between group-1 and group-3 is 2.

To analyze the divergence in specific movie content across developmental stages, we established reference groups at the two extremes: the youngest and the oldest age groups, each comprising 100 individuals. We then calculated the correlation between the local geometry similarity of the remaining individuals and their age, using these two reference groups for comparison. We identified cliques specific to developmental stages, with a significance determined at a threshold of P < 0.05, adjusted for false discovery rate (FDR). Subsequently, we manually annotated those cliques prominently featuring scenes with children or minions (constituting over 50% of screen time). A chi-square test was then performed to determine whether scenes prominently featuring kids or minions significantly contribute to developmental divergence.

### Behavioral association analysis in HBN

For global topology analysis, we divided the participants into five equal-sized groups, ensuring independence in the result. We then calculated the global topology for each participant within these groups ( 5 × 194*subjects* × 750 *volumes*). Subsequently, we computed the Spearman correlation between the principal gradient of the global topology similarity ( 5 × 194*subjects*) and WIAT_LC_Stnd ( 5 × 194*subjects*).

To explore the relationship between local geometry properties and mental health, we implemented two adjustments: (i) considering that psychological issues may not have fully manifested in younger children, we focused our analysis on participants aged above 8 years(*51*), comprising a total of 725 subjects; (ii) to enhance the dynamic specificity of the HBN dataset, we utilized data from both the HCP and a representative subset of the HBN dataset within the STIM framework to fit a hybrid topological space (results presented in Fig. 5i-j). To mitigate the effects of age imbalance, participants were evenly distributed across 10 age groups. The selection of representative HBN time series involved choosing the five volumes with the highest Pearson correlation within each age group at each time point, yielding a dataset with dimensions of 50 × 271 ROIs × 750 volumes. These time series were compiled from multiple individuals, rather than being derived from a single individual source. While clique segmentation was uniform across the individuals, group references were calculated separately for each of the five equal-sized groups based on age ( 5 × 145*subjects*). We standardized the local geometry of individuals within each group and aggregated the data (725*subjects* × 41*cliques*). Following the same process as the HCP data analysis, we performed permutation tests on different cliques to identify probe-like cliques (14 *cliques*) and their paired behavior. Due to the limited number of probe-like cliques identified, we highlighted the top five clique-behavior pairs. Details of all probe-like cliques and their corresponding behaviors with the highest weights are included in Supplementary Tab. 4.

Unlike the HCP data, the fMRI data for HBN were collected from two different sites. Therefore, in all correlation analyses of global topology and local geometry, we controlled for confounding variables by regressing out age, gender, and site information from both the behavioral and topological metrics.

### Statistically testing

To assess statistical significance in our analysis, we adopted the following approach: At the global level, we directly calculated the Spearman correlation between the first principal component of global similarity and various behaviors. Consequently, we conducted FDR cross-validation on 59 trait-like behaviors provided by HCP (Supplementary Tab. 3). In the HBN data, we applied a similar correction for multiple comparisons across different total score items (Supplementary Tab. 3). At the local level, we analyzed the statistical significance between overall behaviors and local geometry. To this end, we generated a null distribution by randomizing local geometry (i.e., shuffling participant IDs) to assess statistical significance. This was done by performing 10,000 random permutations.

## Data availability

The HCP dataset (including neural imaging data, movie stimuli, behavioral measurements and semantic labels) is available at: https://www.humanconnectome.org/study/hcp-young-adult/data-releases. The HBN dataset (including neural imaging data and behavioral measurements) is available at: https://fcon_1000.projects.nitrc.org/indi/cmi_healthy_brain_network/Data.html.

## Code availability

The minimal preprocess pipeline of HCP dataset is freely available at: https://github.com/Washington-University/HCPpipelines. The preprocess of HBN dataset was carried out in fMRIPrep version 20.2.1. fMRIPrep is freely available at: https://github.com/nipreps/fmriprep. Mapper approach was conducted by UAMP version 0.5.3, Kepler Mapper version 1.2.0, and DyNeuSR version 0.3.10. Kepler Mapper is freely available at: https://github.com/scikit-tda/kepler-mapper. UAMP is freely available at: https://github.com/lmcinnes/umap. DyNeuSR is freely available at: https://github.com/braindynamicslab/dyneusr. STIM is available at: https://github.com/JunxingXian/STIM.

## Acknowledgment

This research was conducted using resources from the Human Connectome Project, WU-Minn Consortium (Principal Investigators: David Van Essen and Kamil Ugurbil), and the Healthy Brain Network study (Principal Investigator: Michael P. Milham and Harold S. Koplewicz). We thank Dr. Manish Saggar’s team for developing and maintaining the Mapper analysis tools. We also thank Dr. Danqiu Liu and Dr. Peng Zhang for their valuable advice and assistance during the writing process. This work was supported by STI2030-Major Projects (2022ZD0211902 to A.L.), the Natural Science Foundation of China (grant 82425024 and 82372049 to B.L.; grant 82171543 to A.L.), and the Beijing Nova Program (grant 20230484425 to B.L.).

Author contributions: Conceptualization: A.L., B.L., J.-X.X., Y.-N.H.; Methodology: A.L., B.L., J.-X.X., X.-H.T., Y.-J.P., Y.-N.H., Y.Y., Q.W.; fMRI Data Preprocess: J.-X.X., X.-H.T., Y.-J.P., X.-Y.L., Q.W., P.-Z.L.; Behavioral Data Preprocess: J.-X.X., Y.-N.H., T.G., Y.-J.P.; Data Analysis: J.-X.X., A.L., Y.-N.H., Y.Y., X.-H.T., Y.-J.P., J.L.; Critical Feedback: A.L., B.L., X.-H.T., Y.-J.P., Q.W., Q.W., Y.-Q.S., Y.W., S.-Z. H., K.-X. L., K.H., C.-Y.D., D.-Z.L., M.W.; Writing original draft: J.-X.X., A.L., Y.-N.H., X.-H.T., B.L.; Funding acquisition: A.L., B.L.; Supervision: A.L., B.L.; Competing interests: No competing interests.

## Supplementary Materials

**Fig. S1:**
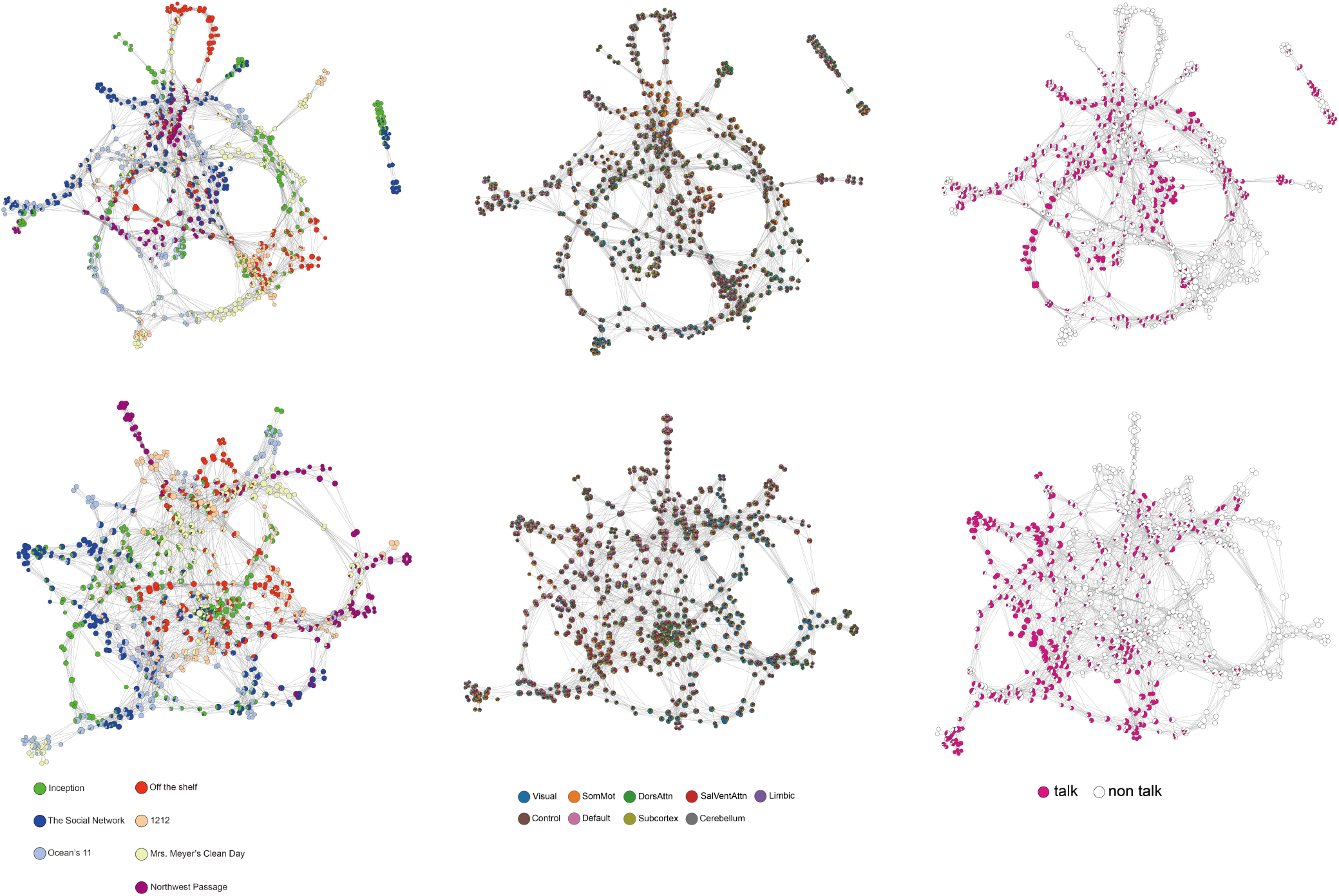
Shape graphs of two participants. **(a)** Two individual shape graphs annotated with movie clips. **(b)** Two individual shape graphs annotated with brain configuration. **(c)** Two individual shape graphs annotated with ‘talk’ semantic label.

**Fig. S2:**
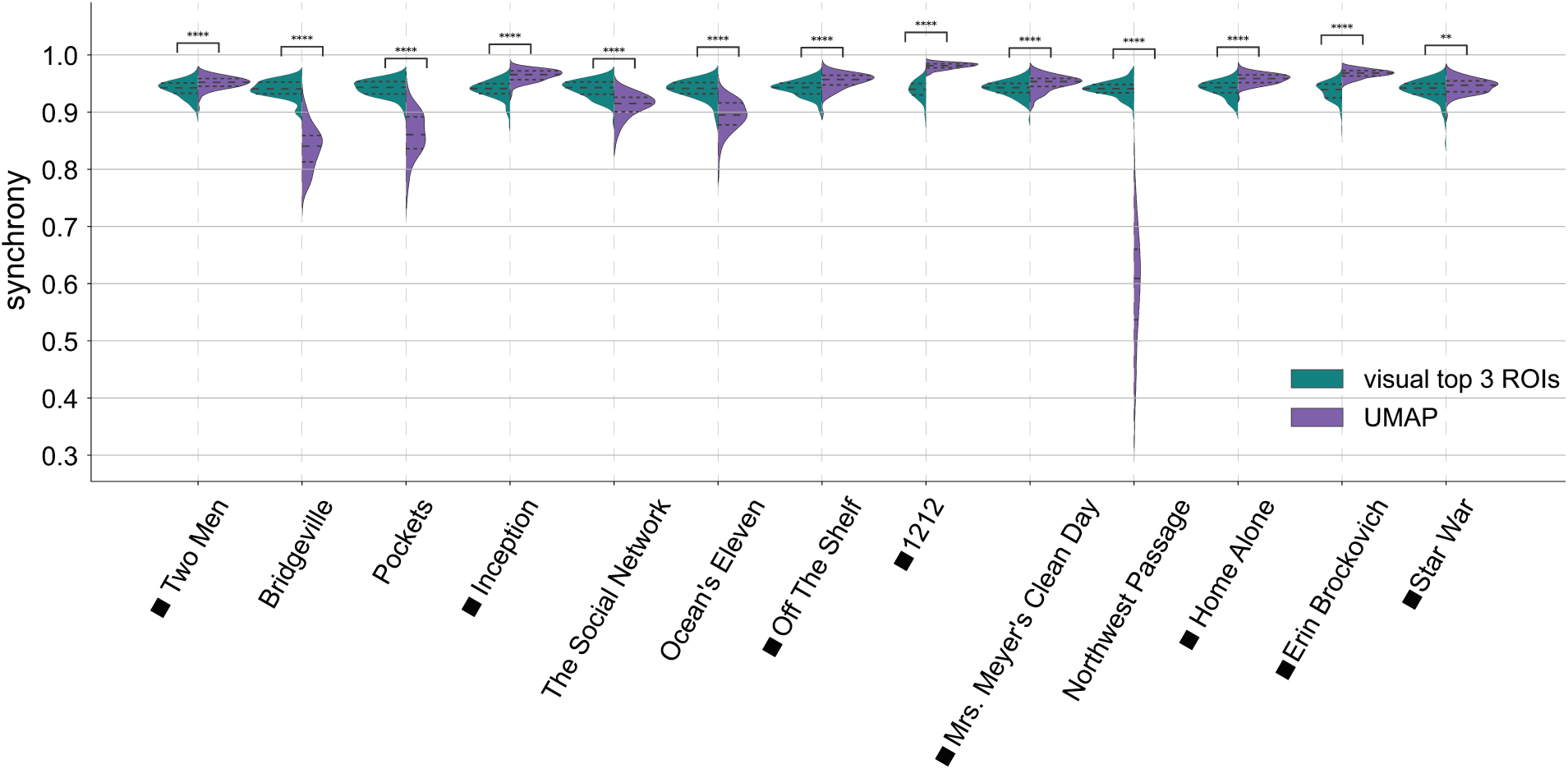
Group synchrony of UMAP embeddings and top visual ROIs. The violin plot illustrates the group synchrony across different movies, comparing UMAP low-dimensional embeddings with the time series of the top three ROIs from the visual network. These top three ROIs were selected based on the highest mean Pearson correlation across all subjects. Synchrony was calculated using the Pearson correlation between two groups, with each group consisting of 15 randomly selected participants, repeated over 100 permutations. The square notation indicates that the UMAP embeddings exhibit significantly higher synchrony compared to the time series data. UMAP embeddings display varying group synchrony across different movie clips, while the time series from the visual network show relatively stable group synchrony. This suggests that the low-dimensional embeddings capture a more personalized cognitive experience of whole-brain integration, rather than merely reflecting visual sensory inputs.

**Fig. S3:**
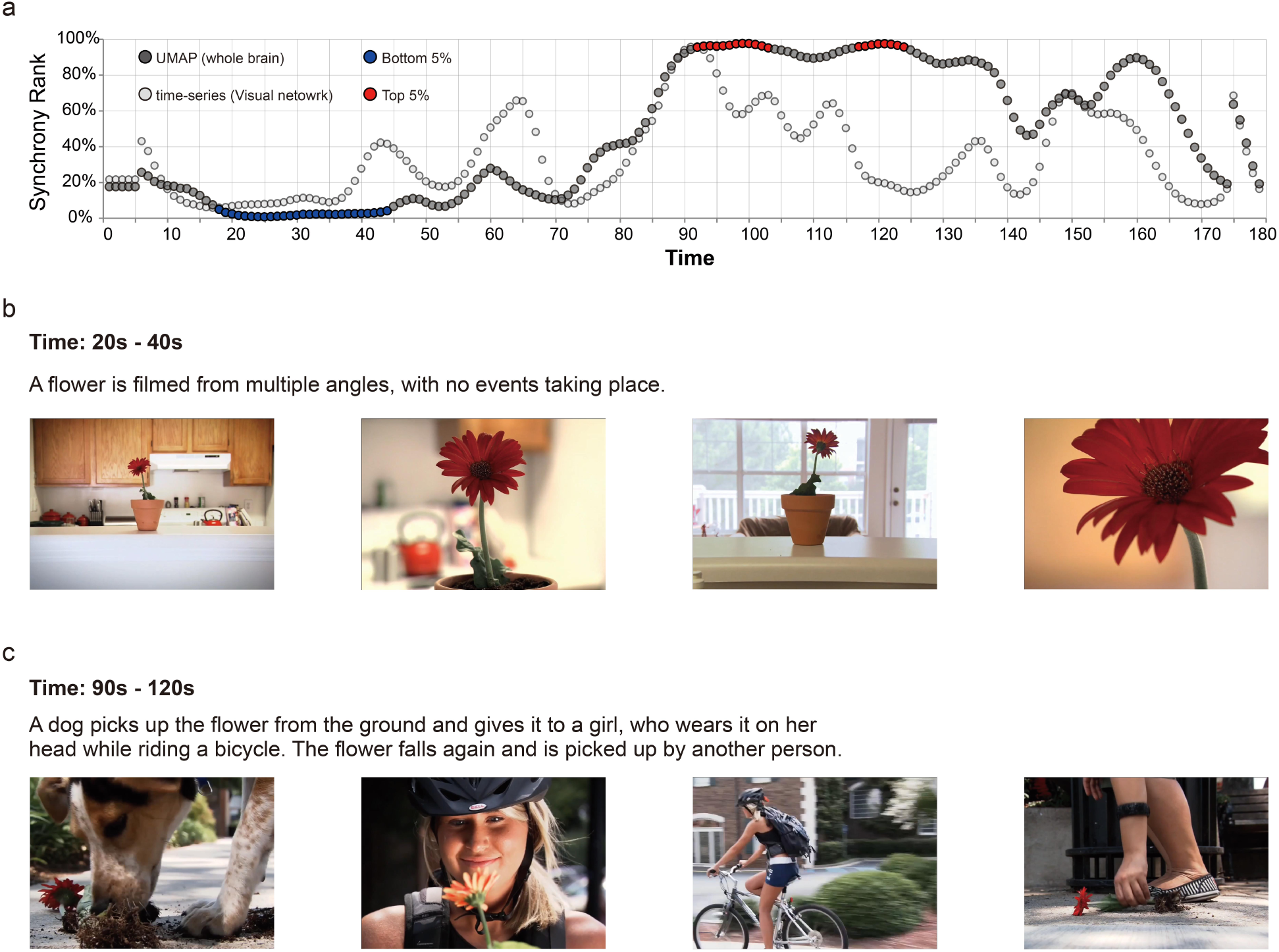
Low-dimensional embeddings integrate subjective experience during movie viewing. To further explore how low-dimensional embeddings of whole-brain dynamics represent subjective experience, we selected a representative clip “Off the Shelf” that contains both non-narrative natural scenes and structured story segments. **(a)** The synchrony rank of whole-brain UMAP embeddings and visual network time-series. Synchrony rank is the rank of the averaged Euclidean distance between all subjects and the group across the entire 13 movie clips. **(b)** Snapshots and text descriptions of the most unconstrained scene (bottom 5%) in “Off the Shelf,” featuring a close-up of a flower. **(c)** Snapshots and text descriptions of the attractor-like scene (top 5%) in “Off the Shelf,” featuring narrative scenes and metaphors.

**Fig. S4:**
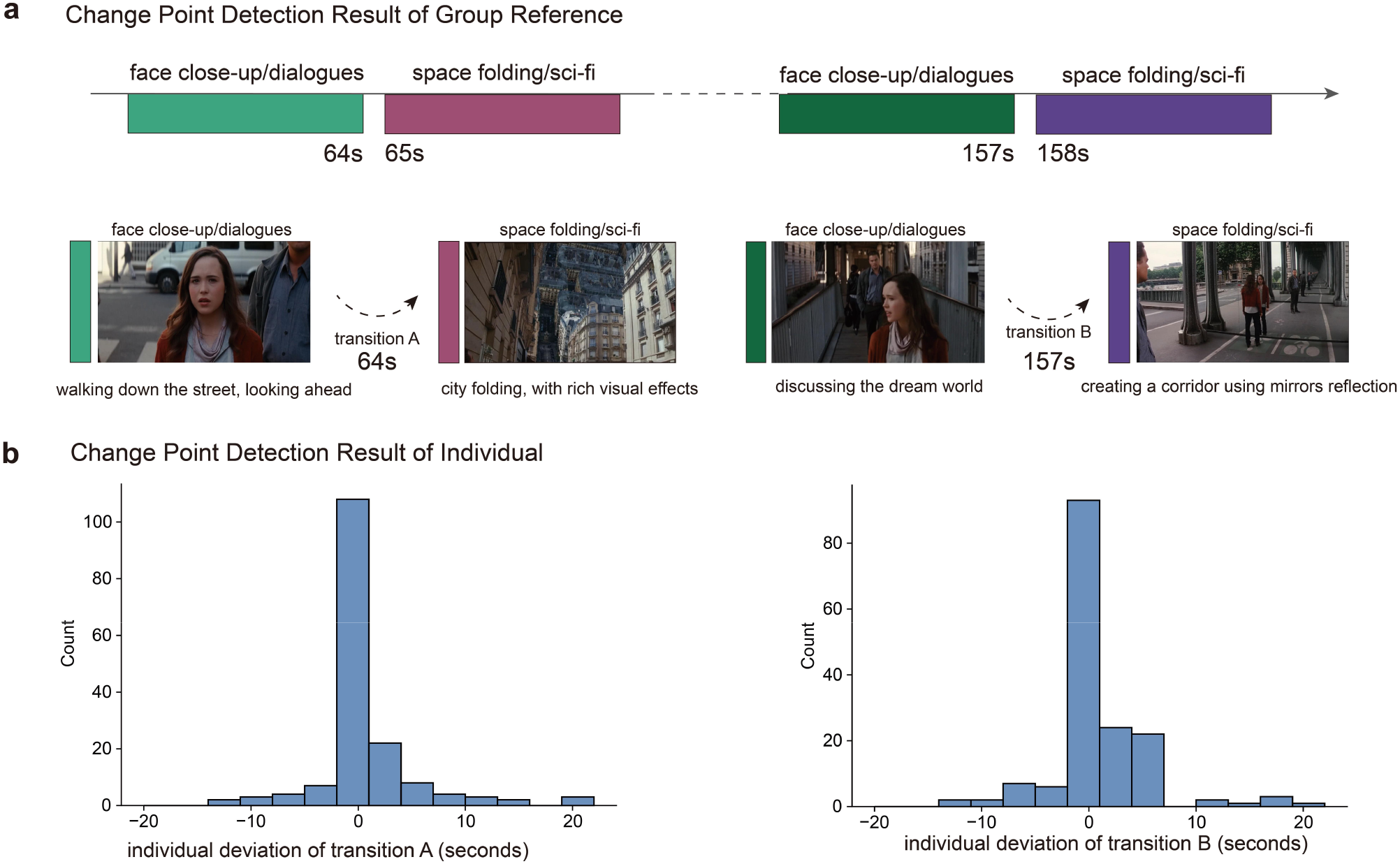
Individual deviation of jumps in *Inception*. This figure displays the deviation of segmentations at jump-A (64 seconds) and jump-B (157 seconds) across all subjects. For example, a subject with a deviation of 2 seconds in jump-A means the clique segmentation of this subject includes 66 seconds.

**Fig. S5:**
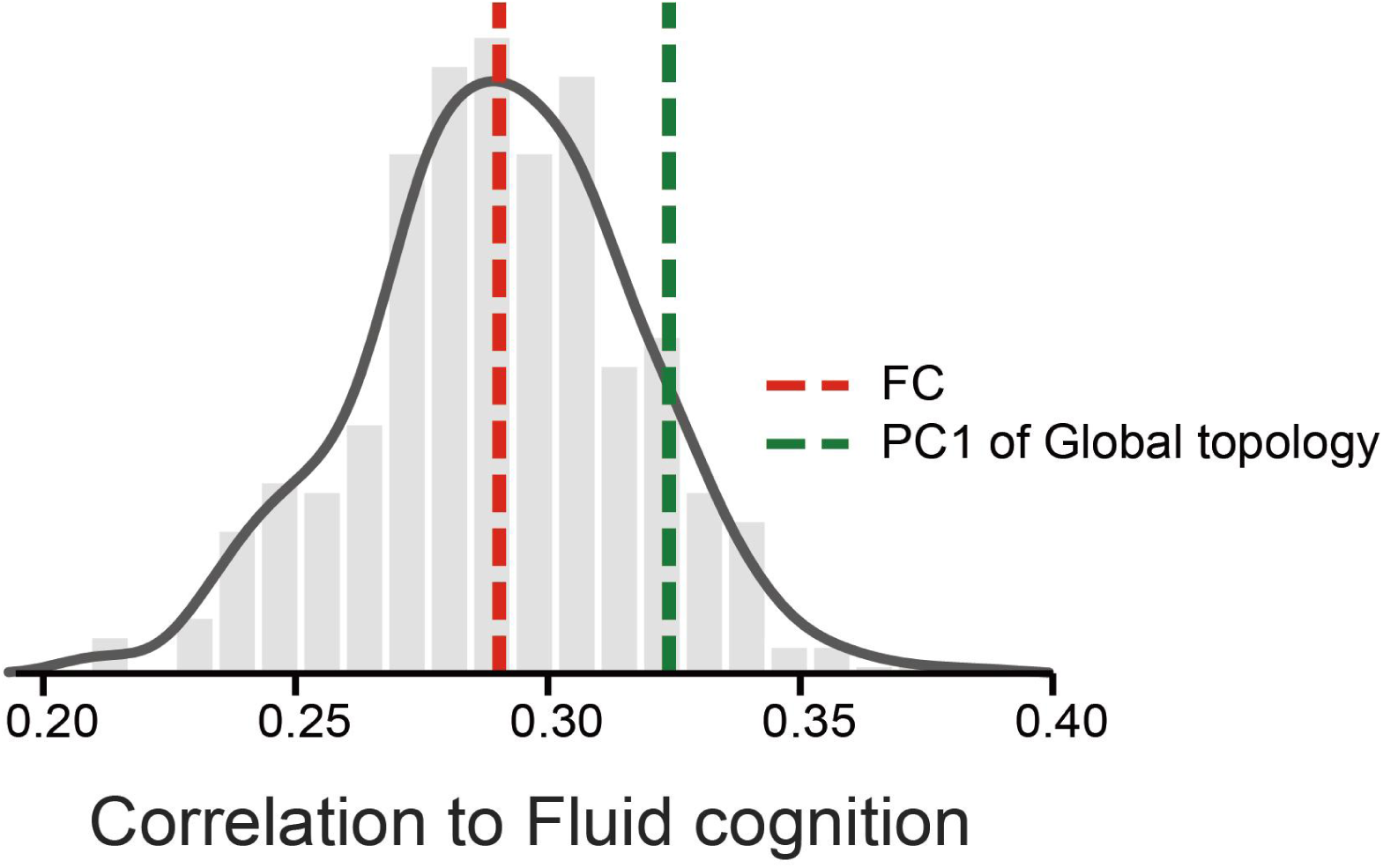
PC1 of global topology captures considerable variance of individual fluid cognition. We computed the Pearson correlation between the time series of different ROIs for each subject, generating a static functional connectome (sFC, dimension: 271*ROIs* × 271*ROIs*). Time series from different movie sessions were concatenated. From this, we extracted the upper triangular matrix as features (dimension: 170 *subjects* × 36856 *features*). An ordinary least squares linear regression model was employed to predict fluid cognition from the sFC, using 5-fold cross-validation. The distribution shown represents the Spearman correlation from 500 different trials, with the mean result marked by a red dotted line. The green dotted line indicates the Spearman correlation between the principal gradient of global topology and fluid cognition. The r-value for global topology is comparable to that of the sFC (P = 0.14), suggesting that PC1 of global topology, a one-dimensional topological measurement, captures a considerable portion of the variance in individual fluid cognition.

**Fig. S6:**
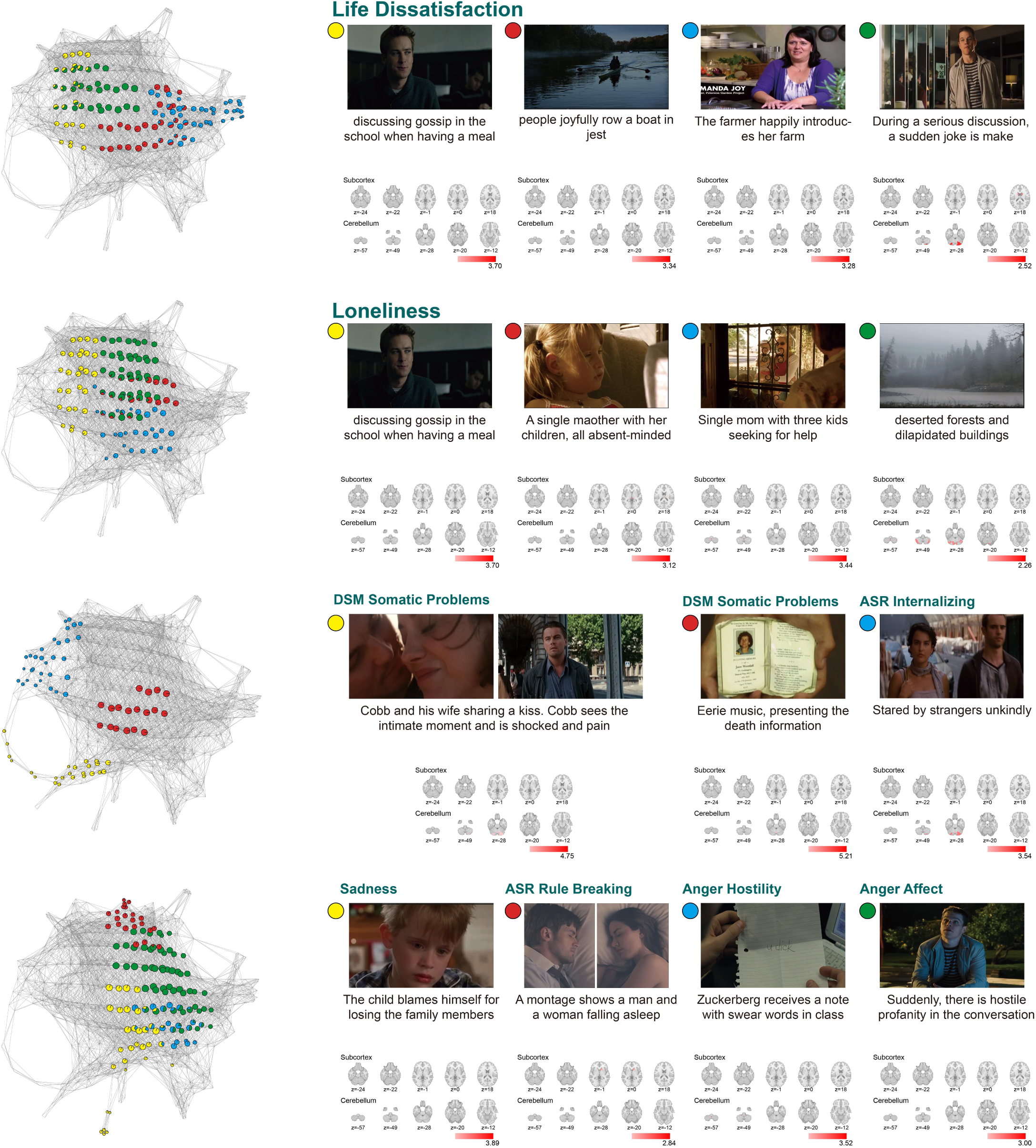
Group network configuration of local geometry. The figure displays the location on the group shape graph of probe-like cliques, showcasing the topological similarity between different cliques. Brain activation maps are z-scored and shown at a threshold of z=1.0 for each probe-like clique.

**Fig. S7:**
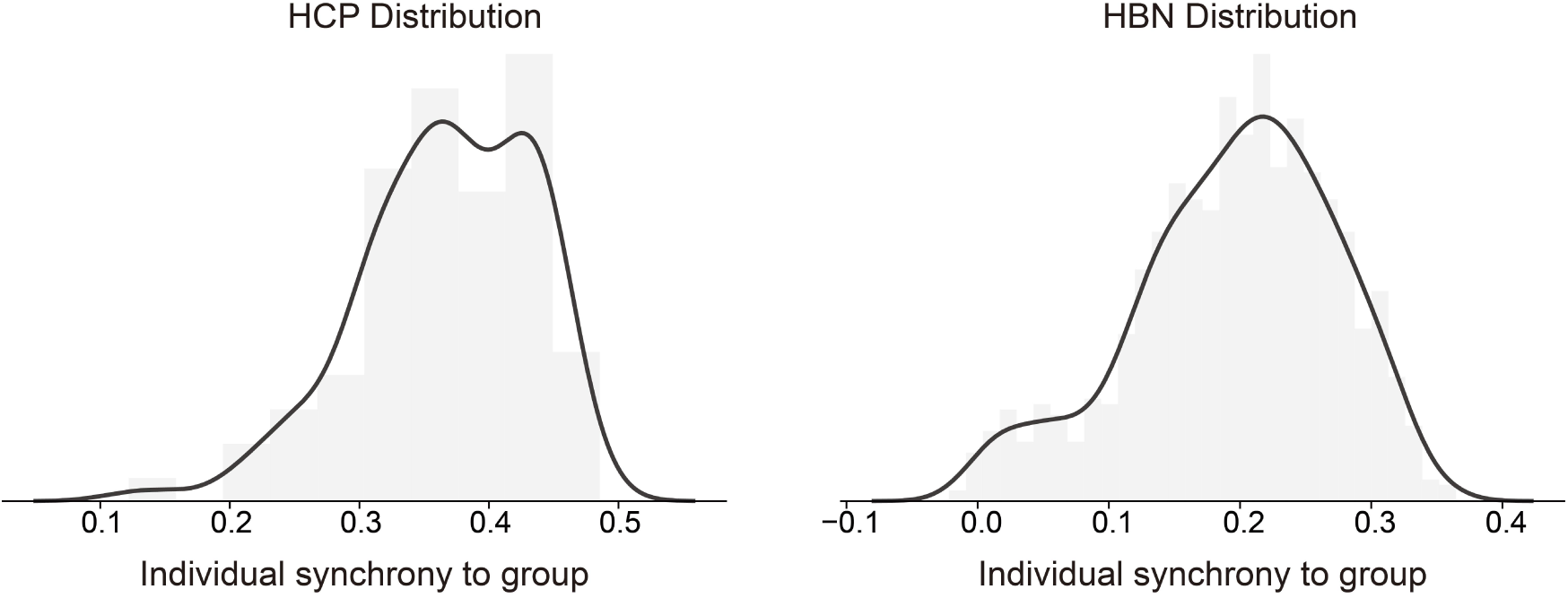
HBN data shows a non-normal distribution of Individual synchrony to group. We examined the individual synchrony to the group in two datasets. The group reference was generated by averaging the time series of all subjects. For each subject, we calculated the Pearson correlation between their time series and the group reference across ROIs. The average of these correlations was used as the measure of individual synchrony. Compared to the HCP data, the HBN data exhibits a non-normal distribution. Left: Distribution of individual synchrony for HCP data. Right: Distribution of individual synchrony for HBN data.

**Fig. S8:**
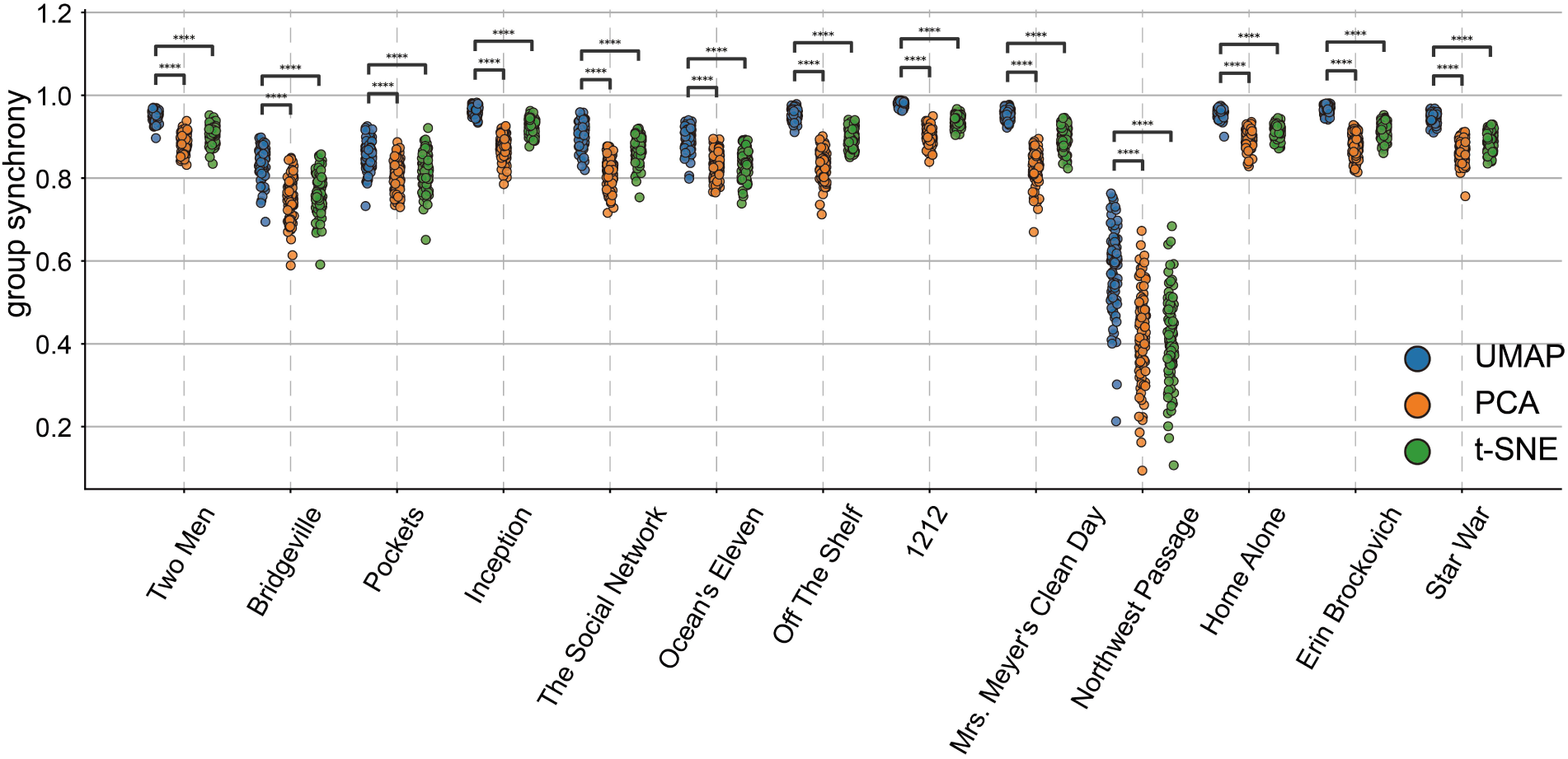
Group synchrony under three different filter functions. The plot shows group synchrony for different movies using three distinct dimensionality reduction techniques: UMAP (blue points), t-SNE (green points), and PCA (orange points). Both UMAP and t-SNE were initialized with PCA. Three filter functions reduce the data to three dimensions. Synchrony was computed as the Pearson correlation between two groups, with each group consisting of 15 randomly selected participants, repeated over 100 permutations.

**Fig. S9:**
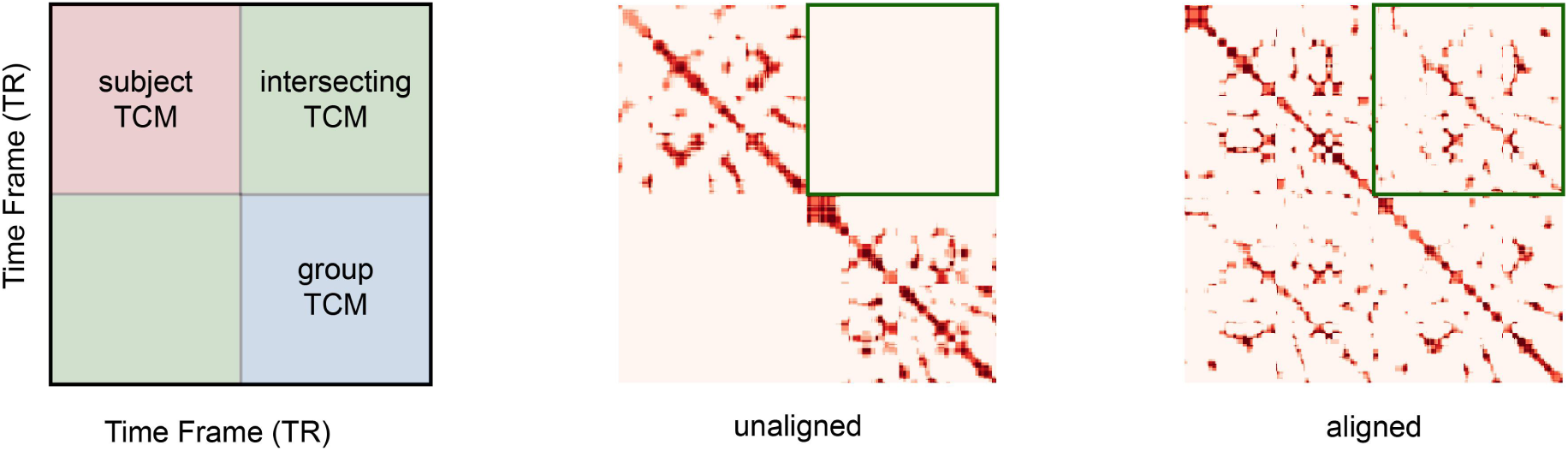
An illustration of intersecting TCM. The illustration shows the intersecting TCM within the STIM framework (in green), highlighting the differences between unaligned and aligned states. The intersecting TCM is used for downstream analysis.

**Fig. S10:**
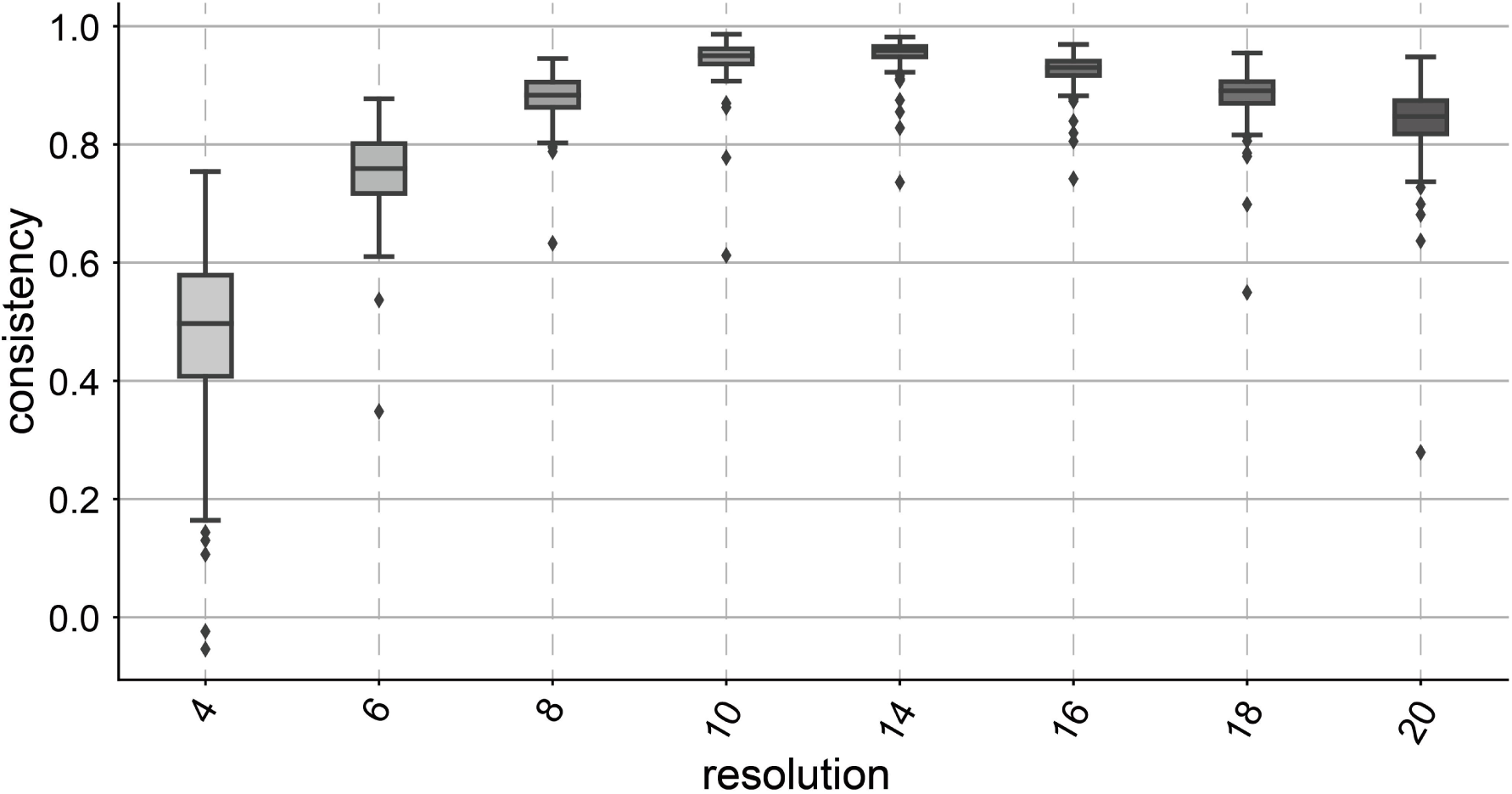
Mapper parameter perturbation test in local geometry. The box plot displays the consistency in local geometry across different resolution parameters, ranging from 4 to 20 (with 12 selected as the reference). Consistency is computed as the Pearson correlation between the test parameter and the reference parameter (resolution = 12).

## Reference

1. D. M. Barch, G. C. Burgess, M. P. Harms, S. E. Petersen, B. L. Schlaggar, M. Corbetta, M. F. Glasser, S. Curtiss, S. Dixit, C. Feldt, D. Nolan, E. Bryant, T. Hartley, O. Footer, J. M. Bjork, R. Poldrack, S. Smith, H. Johansen-Berg, A. Z. Snyder, D. C. Van Essen, WU-Minn HCP Consortium, Function in the human connectome: task-fMRI and individual differences in behavior. Neuroimage 80, 169–189 (2013).

2. J. Gonzalez-Castillo, C. W. Hoy, D. A. Handwerker, M. E. Robinson, L. C. Buchanan, Z. S. Saad, P. A. Bandettini, Tracking ongoing cognition in individuals using brief, whole-brain functional connectivity patterns. Proc Natl Acad Sci U S A 112, 8762–8767 (2015).

3. W. Zhao, C. Makowski, D. J. Hagler, H. P. Garavan, W. K. Thompson, D. J. Greene, T. L. Jernigan, A. M. Dale, Task fMRI paradigms may capture more behaviorally relevant information than resting-state functional connectivity. Neuroimage 270, 119946 (2023).

4. R. Liégeois, J. Li, R. Kong, C. Orban, D. Van De Ville, T. Ge, M. R. Sabuncu, B. T. T. Yeo, Resting brain dynamics at different timescales capture distinct aspects of human behavior. Nat Commun 10, 1–9 (2019).

5. E. Tagliazucchi, H. Laufs, Decoding Wakefulness Levels from Typical fMRI Resting-State Data Reveals Reliable Drifts between Wakefulness and Sleep. Neuron 82, 695–708 (2014).

6. A. Li, H. Liu, X. Lei, Y. He, Q. Wu, Y. Yan, X. Zhou, X. Tian, Y. Peng, S. Huang, K. Li, M. Wang, Y. Sun, H. Yan, C. Zhang, S. He, R. Han, X. Wang, B. Liu, Hierarchical fluctuation shapes a dynamic flow linked to states of consciousness. Nat Commun 14, 3238 (2023).

7. E. S. Finn, P. A. Bandettini, Movie-watching outperforms rest for functional connectivity-based prediction of behavior. Neuroimage 235, 117963 (2021).

8. S. Sonkusare, M. Breakspear, C. Guo, Naturalistic Stimuli in Neuroscience: Critically Acclaimed. Trends Cogn Sci 23, 699–714 (2019).

9. I. P. Jääskeläinen, M. Sams, E. Glerean, J. Ahveninen, Movies and narratives as naturalistic stimuli in neuroimaging. Neuroimage 224, 117445 (2021).

10. U. Hasson, Y. Nir, I. Levy, G. Fuhrmann, R. Malach, Intersubject synchronization of cortical activity during natural vision. Science 303, 1634–1640 (2004).

11. J.-P. Kauppi, I. P. Jääskeläinen, M. Sams, J. Tohka, Inter-subject correlation of brain hemodynamic responses during watching a movie: localization in space and frequency. Front. Neuroinform. 4, 669 (2010).

12. S. A. Nastase, V. Gazzola, U. Hasson, C. Keysers, Measuring shared responses across subjects using intersubject correlation. Soc Cogn Affect Neurosci 14, 667–685 (2019).

13. E. S. Finn, E. Glerean, A. Y. Khojandi, D. Nielson, P. J. Molfese, D. A. Handwerker, P. A. Bandettini, Idiosynchrony: From shared responses to individual differences during naturalistic neuroimaging. Neuroimage 215, 116828 (2020).

14. P.-H. A. Chen, E. Jolly, J. H. Cheong, L. J. Chang, Intersubject representational similarity analysis reveals individual variations in affective experience when watching erotic movies. Neuroimage 216, 116851 (2020).

15. E. Simony, C. J. Honey, J. Chen, O. Lositsky, Y. Yeshurun, A. Wiesel, U. Hasson, Dynamic reconfiguration of the default mode network during narrative comprehension. Nat Commun 7, 1–13 (2016).

16. J. N. van der Meer, M. Breakspear, L. J. Chang, S. Sonkusare, L. Cocchi, Movie viewing elicits rich and reliable brain state dynamics. Nat Commun 11, 1–14 (2020).

17. H. Song, W. M. Shim, M. D. Rosenberg, Large-scale neural dynamics in a shared low-dimensional state space reflect cognitive and attentional dynamics, eLife (2023). 10.7554/eLife.85487.

18. L. G. A. Freitas, T. A. W. Bolton, B. E. Krikler, D. Jochaut, A.-L. Giraud, P. S. Hüppi, D. Van De Ville, Time-resolved effective connectivity in task fMRI: Psychophysiological interactions of Co-Activation patterns. Neuroimage 212, 116635 (2020).

19. J. M. Shine, M. Breakspear, P. T. Bell, K. A. Ehgoetz Martens, R. Shine, O. Koyejo, O. Sporns, R. A. Poldrack, Human cognition involves the dynamic integration of neural activity and neuromodulatory systems. Nat Neurosci 22, 289–296 (2019).

20. R. Chaudhuri, B. Gerçek, B. Pandey, A. Peyrache, I. Fiete, The intrinsic attractor manifold and population dynamics of a canonical cognitive circuit across waking and sleep. Nat Neurosci 22, 1512–1520 (2019).

21. S. S. Kim, H. Rouault, S. Druckmann, V. Jayaraman, Ring attractor dynamics in the Drosophila central brain. Science 356, 849–853 (2017).

22. R. J. Gardner, E. Hermansen, M. Pachitariu, Y. Burak, N. A. Baas, B. A. Dunn, M.-B. Moser, E. I. Moser, Toroidal topology of population activity in grid cells. Nature 602, 123–128 (2022).

23. G. Singh, F. Memoli, G. Carlsson, Topological Methods for the Analysis of High Dimensional Data Sets and 3D Object Recognition (The Eurographics Association, 2007; 10.2312/SPBG/SPBG07/091-100).

24. M. Saggar, O. Sporns, J. Gonzalez-Castillo, P. A. Bandettini, G. Carlsson, G. Glover, A. L. Reiss, Towards a new approach to reveal dynamical organization of the brain using topological data analysis. Nat Commun 9, 1–14 (2018).

25. M. Saggar, J. M. Shine, R. Liégeois, N. U. F. Dosenbach, D. Fair, Precision dynamical mapping using topological data analysis reveals a hub-like transition state at rest. Nat Commun 13, 1–19 (2022).

26. C. Geniesse, O. Sporns, G. Petri, M. Saggar, Generating dynamical neuroimaging spatiotemporal representations (DyNeuSR) using topological data analysis. Network Neuroscience 3, 763 (2019).

27. L. McInnes, J. Healy, J. Melville, UMAP: Uniform Manifold Approximation and Projection for Dimension Reduction, arXiv.org (2018). https://arxiv.org/abs/1802.03426v3.

28. L. van der Maaten, G. Hinton, Visualizing Data using t-SNE. Journal of Machine Learning Research 9, 2579–2605 (2008).

29. J. Tseng, J. Poppenk, Brain meta-state transitions demarcate thoughts across task contexts exposing the mental noise of trait neuroticism. Nat Commun 11, 3480 (2020).

30. E. C. Baek, R. Hyon, K. López, E. S. Finn, M. A. Porter, C. Parkinson, In-degree centrality in a social network is linked to coordinated neural activity. Nat Commun 13, 1118 (2022).

31. R. Killick, P. Fearnhead, I. A. Eckley, Optimal Detection of Changepoints With a Linear Computational Cost. Journal of the American Statistical Association 107, 1590–1598 (2012).

32. L. C. Freeman, A set of measures of centrality based on betweenness. Sociometry, 35–41 (1977).

33. J. F. Brosschot, Cognitive-emotional sensitization and somatic health complaints. Scandinavian journal of psychology 43 (2002).

34. Y. Zhang, J.-H. Kim, D. Brang, Z. Liu, Naturalistic Stimuli: A Paradigm for Multi-Scale Functional Characterization of the Human Brain. Current opinion in biomedical engineering 19 (2021).

35. S. B. Eickhoff, M. Milham, T. Vanderwal, Towards clinical applications of movie fMRI. Neuroimage 217, 116860 (2020).

36. X. Li, S. B. Eickhoff, S. Weis, Stimulus selection influences prediction of individual phenotypes in naturalistic conditions. bioRxiv, 2023.12.07.570273 (2023).

37. L. J. Chang, E. Jolly, J. H. Cheong, K. M. Rapuano, N. Greenstein, P.-H. A. Chen, J. R. Manning, Endogenous variation in ventromedial prefrontal cortex state dynamics during naturalistic viewing reflects affective experience. Science Advances, doi: 10.1126/sciadv.abf7129 (2021).

38. E. Morgenroth, L. Vilaclara, M. Muszynski, J. Gaviria, P. Vuilleumier, D. Van De Ville, Probing neurodynamics of experienced emotions—a Hitchhiker’s guide to film fMRI. Soc Cogn Affect Neurosci 18 (2023).

39. G. S. Howard, Response-Shift Bias. Evaluation Review, doi: 10.1177/0193841X8000400105 (1980).

40. K. M. Lyons, R. A. Stevenson, A. M. Owen, B. Stojanoski, Examining the relationship between measures of autistic traits and neural synchrony during movies in children with and without autism. Neuroimage Clin 28, 102477 (2020).

41. D. C. Gruskin, M. D. Rosenberg, A. J. Holmes, Relationships between depressive symptoms and brain responses during emotional movie viewing emerge in adolescence. Neuroimage 216, 116217 (2020).

42. Z. Yang, J. Wu, L. Xu, Z. Deng, Y. Tang, J. Gao, Y. Hu, Y. Zhang, S. Qin, C. Li, J. Wang, Individualized psychiatric imaging based on inter-subject neural synchronization in movie watching. Neuroimage 216, 116227 (2020).

43. S. Schneider, J. H. Lee, M. W. Mathis, Learnable latent embeddings for joint behavioural and neural analysis. Nature 617, 360–368 (2023).

44. C. Geniesse, S. Chowdhury, M. Saggar, NeuMapper: A scalable computational framework for multiscale exploration of the brain’s dynamical organization. Netw Neurosci 6, 467–498 (2022).

45. M. C. Camacho, A. N. Nielsen, D. Balser, E. Furtado, D. C. Steinberger, L. Fruchtman, J. P. Culver, C. M. Sylvester, D. M. Barch, Large-scale encoding of emotion concepts becomes increasingly similar between individuals from childhood to adolescence. Nat Neurosci 26, 1256–1266 (2023).

46. L. Nummenmaa, E. Glerean, M. Viinikainen, I. P. Jääskeläinen, R. Hari, M. Sams, Emotions promote social interaction by synchronizing brain activity across individuals. doi: 10.1073/pnas.1206095109 (2012).

47. D. C. Van Essen, S. M. Smith, D. M. Barch, T. E. J. Behrens, E. Yacoub, K. Ugurbil, WU-Minn HCP Consortium, The WU-Minn Human Connectome Project: an overview. Neuroimage 80, 62–79 (2013).

48. D. C. Van Essen, K. Ugurbil, E. Auerbach, D. Barch, T. E. J. Behrens, R. Bucholz, A. Chang, L. Chen, M. Corbetta, S. W. Curtiss, S. Della Penna, D. Feinberg, M. F. Glasser, N. Harel, A. C. Heath, L. Larson-Prior, D. Marcus, G. Michalareas, S. Moeller, R. Oostenveld, S. E. Petersen, F. Prior, B. L. Schlaggar, S. M. Smith, A. Z. Snyder, J. Xu, E. Yacoub, WU-Minn HCP Consortium, The Human Connectome Project: a data acquisition perspective. Neuroimage 62, 2222–2231 (2012).

49. M. F. Glasser, S. N. Sotiropoulos, J. A. Wilson, T. S. Coalson, B. Fischl, J. L. Andersson, J. Xu, S. Jbabdi, M. Webster, J. R. Polimeni, D. C. Van Essen, M. Jenkinson, WU-Minn HCP Consortium, The minimal preprocessing pipelines for the Human Connectome Project. Neuroimage 80, 105–124 (2013).

50. K. J. Friston, S. Williams, R. Howard, R. S. Frackowiak, R. Turner, Movement-related effects in fMRI time-series. Magn Reson Med 35, 346–355 (1996).

51. L. M. Alexander, J. Escalera, L. Ai, C. Andreotti, K. Febre, A. Mangone, N. Vega-Potler, N. Langer, A. Alexander, M. Kovacs, S. Litke, B. O’Hagan, J. Andersen, B. Bronstein, A. Bui, M. Bushey, H. Butler, V. Castagna, N. Camacho, E. Chan, D. Citera, J. Clucas, S. Cohen, S. Dufek, M. Eaves, B. Fradera, J. Gardner, N. Grant-Villegas, G. Green, C. Gregory, E. Hart, S. Harris, M. Horton, D. Kahn, K. Kabotyanski, B. Karmel, S. P. Kelly, K. Kleinman, B. Koo, E. Kramer, E. Lennon, C. Lord, G. Mantello, A. Margolis, K. R. Merikangas, J. Milham, G. Minniti, R. Neuhaus, A. Levine, Y. Osman, L. C. Parra, K. R. Pugh, A. Racanello, A. Restrepo, T. Saltzman, B. Septimus, R. Tobe, R. Waltz, A. Williams, A. Yeo, F. X. Castellanos, A. Klein, T. Paus, B. L. Leventhal, R. C. Craddock, H. S. Koplewicz, M. P. Milham, An open resource for transdiagnostic research in pediatric mental health and learning disorders. Sci Data 4, 1–26 (2017).

52. O. Esteban, C. J. Markiewicz, R. W. Blair, C. A. Moodie, A. I. Isik, A. Erramuzpe, J. D. Kent, M. Goncalves, E. DuPre, M. Snyder, H. Oya, S. S. Ghosh, J. Wright, J. Durnez, R. A. Poldrack, K. J. Gorgolewski, fMRIPrep: a robust preprocessing pipeline for functional MRI. Nat Methods 16, 111–116 (2019).

53. B. Liu, X. Tian, Y. Peng, M. Wang, Y. Sun, J. Lou, J. Xian, Y. He, K. Hu, Q. Wang, S. Huang, A. Li, Deciphering Complex Brain Spatiotemporal Dynamics Shaping Diverse Human Behavior (2023).

54. A. Schaefer, R. Kong, E. M. Gordon, T. O. Laumann, X.-N. Zuo, A. J. Holmes, S. B. Eickhoff, B. T. T. Yeo, Local-Global Parcellation of the Human Cerebral Cortex from Intrinsic Functional Connectivity MRI. Cereb Cortex 28, 3095–3114 (2018).

55. Y. Tian, D. S. Margulies, M. Breakspear, A. Zalesky, Topographic organization of the human subcortex unveiled with functional connectivity gradients. Nat Neurosci 23, 1421–1432 (2020).

56. R. L. Buckner, F. M. Krienen, A. Castellanos, J. C. Diaz, B. T. T. Yeo, The organization of the human cerebellum estimated by intrinsic functional connectivity. J Neurophysiol 106, 2322–2345 (2011).

57. H. Lee, J. Chen, Predicting memory from the network structure of naturalistic events. Nat Commun 13, 4235 (2022).

58. H. J. van Veen, N. Saul, D. Eargle, S. W. Mangham, Kepler Mapper: A flexible Python implementation of the Mapper algorithm. Journal of Open Source Software 4, 1315 (2019).

59. M. Ester, H.-P. Kriegel, J. Sander, X. Xu, “A density-based algorithm for discovering clusters in large spatial databases with noise” in Proceedings of the Second International Conference on Knowledge Discovery and Data Mining (AAAI Press, Portland, Oregon, 1996)KDD’96, pp. 226–231.

60. E. L. Busch, J. Huang, A. Benz, T. Wallenstein, G. Lajoie, G. Wolf, S. Krishnaswamy, N. B. Turk-Browne, Multi-view manifold learning of human brain-state trajectories. Nat Comput Sci 3, 240–253 (2023).

61. D. F. Crouse, On implementing 2D rectangular assignment algorithms. IEEE Transactions on Aerospace and Electronic Systems 52, 1679–1696 (2016).

62. D. Haşegan, C. Geniesse, S. Chowdhury, M. Saggar, Deconstructing the Mapper algorithm to extract richer topological and temporal features from functional neuroimaging data. Network Neuroscience, 1–60 (2024).

63. W. B. Bilker, J. A. Hansen, C. M. Brensinger, J. Richard, R. E. Gur, R. C. Gur, Development of abbreviated nine-item forms of the Raven’s standard progressive matrices test. Assessment 19, 354–369 (2012).

